# MKMC enables reference-free transcriptomic analysis using k-mer representations

**DOI:** 10.64898/2026.07.06.736868

**Authors:** Lajoyce Mboning, Maciej D-lugosz, Marek Kokot, Jingxun Chen, Emma K. Costa, Man-Ru Wu, Sui Wang, Louis-S Bouchard, Sebastian Deorowicz, Matteo Pellegrini

## Abstract

Traditional RNA-seq analysis depends heavily on genome alignment and gene annotation, limiting its utility in non-model organisms and introducing biases that can obscure regulatory complexity. We present MKMC (Multi-sample Kmer Counter), a scalable, reference-free toolkit for RNA-seq analysis that leverages k-mer–based statistics to detect biological variation without requiring alignment. MKMC integrates fast k-mer counting, abundance matrix generation, normalization, dimensionality reduction, and differential analysis into a unified workflow. Across diverse datasets, MKMC recapitulates key biological signals—including sex differences in killifish liver—and matches alignment-based pipelines in differential expression analysis and transcriptomic age prediction. Notably, MKMC detects isoform-specific events missed by traditional methods, one of which we validated using in situ hybridization. These results reveal previously hidden isoform-level regulatory events that contribute to sex-and age-associated transcriptional programs. MKMC offers a robust, extensible alternative to alignment-based approaches, enabling transcriptomic discovery across both model and non-model systems. While we focus here on RNA-seq as a primary application, MKMC is broadly applicable to any k-mer–based analysis of next-generation sequencing data.

## 1 Main

High-throughput sequencing technologies generate vast amounts of data across diverse modalities, including genomics, transcriptomics, and epigenomics. A fundamental strategy for analyzing such data is the use of k-mer–based approaches, which decompose sequences into short, fixed-length substrings (k-mers) to enable efficient, alignment-free analysis [1]. By operating directly on raw sequencing reads, k-mer methods provide an annotation-agnostic framework that facilitates scalable quantification, pattern discovery, and the identification of novel sequence features without reliance on reference genomes or transcript models[2, 3]. Despite their conceptual advantages, existing k-mer–based tools often face limitations in scalability, flexibility, and usability[4–6]. Many lack support for multi-condition comparisons, efficient construction of abundance matrices across samples, or seamless integration with downstream statistical and machine learning workflows, limiting their utility in large-scale biological studies[1, 7]. RNA sequencing (RNA-seq) has revolutionized transcriptomic analysis by enabling genome-wide quantification of gene expression across diverse biological conditions and species[8]. However, traditional RNA-seq analysis relies heavily on alignment-based pipelines that map sequencing reads to a reference genome or transcriptome, followed by quantification of gene-or transcript-level abundance[9–12]. While effective in well-annotated model organisms, these approaches face key challenges in species with incomplete genomes and in the discovery of unannotated or novel transcripts[4–6]. A central limitation of alignment-based methods is their dependency on existing annotations, which can obscure biologically meaningful signals arising from novel transcripts, unannotated isoforms, or genomic regions with structural variation [4–6]. Additionally, alignment introduces biases through parameter choices, heuristic filtering, and reliance on transcript models, potentially limiting sensitivity in noisy or highly dynamic biological contexts[2, 3]. To address some of these challenges, we developed MKMC (Multi-sample K-mer Counter), a scalable, reference-free toolkit for k-mer–based analysis of next-generation sequencing data. While broadly applicable across sequencing modalities, we focus here on RNA-seq as a representative use case. MKMC integrates fast k-mer counting, abundance matrix construction, differential k-mer discovery across multiple conditions, and downstream analysis with machine learning into a unified workflow. Designed for speed, modularity, and accessibility, MKMC supports both exploratory analyses and hypothesis-driven investigations in model and non-model organisms alike. In this paper, we describe the design and capabilities of MKMC, benchmark its performance against existing tools, and demonstrate its utility across biological case studies, including differential expression analysis and transcriptomic age prediction.

## 2 MKMC workflow

### 2.1 Workflow overview

The Multi-sample *k*-mer counter (MKMC) is a versatile tool for alignment-free analysis of sequencing datasets. While we use RNA-seq data as a primary example throughout this study, the framework is broadly applicable to other sequencing modalities. MKMC is designed as a multi-stage application (Fig. 1), though all stages can be executed with a single command.

**Fig. 1:**
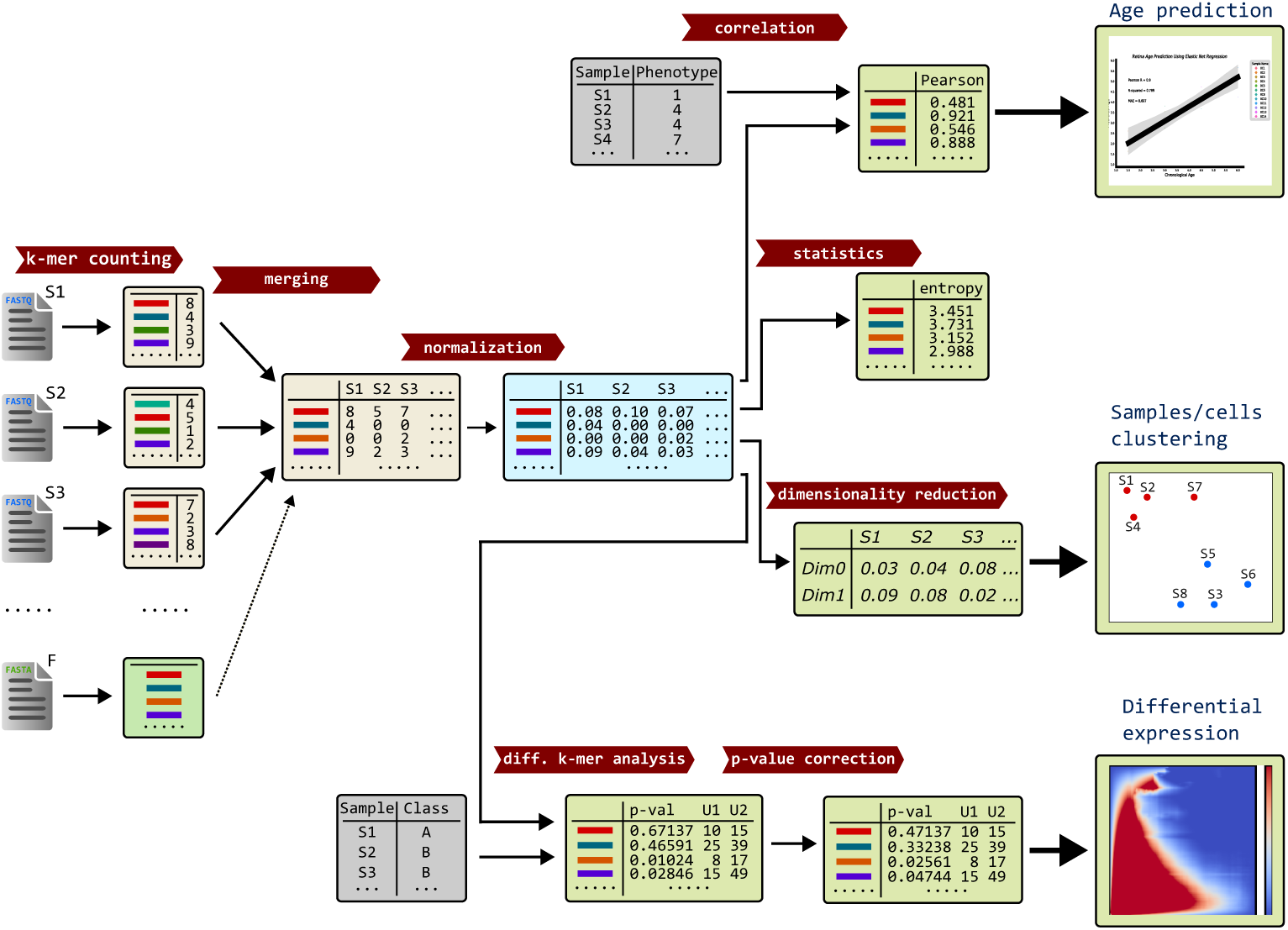
MKMC pipeline and main features. Illustration of the MKMC pipeline. The input is a set of sample data in FASTQ format (S1,…). Optionally, the user can pass a FASTA/text file *F* for filtering. After counting *k*-mers and merging the results, a single matrix *M* is obtained. The counters in *M* are usually normalized (several algorithms implemented). Then, the user can select one or more scenarios for data analysis. The figure illustrates just a few examples (e.g., Pearson correlation), but many more are implemented (details are provided in the main text).

The first stage involves counting *k*-mers separately in all input samples. We use a modified KMC [13] utility here. Then, in the merging stage, *k*-mers with their counts in different samples are merged to construct a matrix, where each row represents a unique *k*-mer and each column represents a sample, with entries containing the count of a given *k*-mer in a specific sample. This stage also collects data for further normalization, and, optionally, generates the output text matrix and FASTA file. During merging, various filtering criteria can be applied (optionally), such as removing *k*-mers that occur less frequently than a certain threshold in a given fraction of samples, or removing *k*-mers absent from a user-provided list.

The following stages depend on the user-selected work scenario (many of them can be selected simultaneously). In all of them, MKMC can postprocess normalized counts or non-normalized (depending on the user’s needs). First, we can calculate the correlation with the provided list of numeric values, one per sample (e.g., phenotypes). Optionally, we can use cross-validation here. Second, we can perform a differential *k*-mer analysis using a provided list of identifiers (labels or numbers, one per sample). The obtained *p*-values can be corrected by one of the provided methods. Third, we can calculate some statistics. Currently, only entropy calculation is implemented; however, the list will be expanded in the future. Fourth, we can perform dimensionality reduction using PCA or UMAP methods. The results of the analyses are stored in separate text files.

MKMC is implemented in the C++ 20 programming language. The application is highly parallelized to utilize modern multi-core CPUs. Nevertheless, there are parts where we employ external programming libraries (e.g., for dimensionality reduction), and some of these libraries are single-threaded.

### 2.2 *k*-mer counting

The *k*-mer counting stage is performed by running KMC for each sample separately, generating a single KMCDB file per sample. The *k*-mers are distributed into (by default 512; every KMC instance is configured to have the same number) containers called *bins*. They are units of data decomposition in later parallel processing. This is made possible by a KMC modification, ensuring that the specific *k*-mer will always land in the bin of the same ID for each sample in which it occurs. This ensures that bins can be processed independently in later stages, each by a separate thread. Another rationale for storing *k*-mers in bins is the concept of KMC flow, as explained in [13]. The original KMC was modified by introducing KMCDB format (see Section 2.5), changing the *k*-mers signature computation method to MinHash, and updating the method of assigning signatures to bin IDs, ensuring the files’ compatibility (see Section 2.5). MKMC is responsible for parameterising it, running it, and later utilizing the generated files.

The user can optionally pass a set *F* of *k*-mers (as a text file or a FASTA file; in particular, *F* may be generated by extracting *k*-mers from, e.g., a genome), which are the only ones that should be considered (other *k*-mers will be filtered out in a merging stage). If so, they are also counted by KMC in the aim of obtaining a compatible KMCDB file.

The user can also provide some filtering criteria at this stage. For example, *k*-mers that are less frequent or more frequent than the given thresholds can be removed during counting. Especially setting the lower threshold to more than 1 (default) could remarkably improve the running time of *k*-mer counting as well as reduce memory and disk usage.

Multiple instances of KMC are run simultaneously. KMC itself is a parallel algorithm; however, since a single sample typically consists of only one or two gzipped FASTQ files, it is often the case that decompression is a bottleneck, limiting throughput. Thus, simultaneous KMC execution allows for better thread utilization. On the other hand, running too many instances may harm disk access patterns (especially in the case of HDD) and increase RAM requirements. For this reason, as a compromise, we have chosen four as the default number of KMC instances to run simultaneously.

### 2.3 Merging samples

The input of the merging stage is a sequence of *n* KMCDB files (one per sample) with *k*-mers and their counts in each of the samples. The aim is to build a matrix *M* with *n* columns. Rows will contain the number of occurrences of each unique *k*-mer *x* from the input samples in each of the samples: *c*(*x, j*), for *j* being the sample ID (although some filtering can be applied; discussed below). The rows of the matrix are labeled by *k*-mers and the columns by samples.

To merge all *k*-mers across samples for a given bin, a binary heap of size *n* is employed, where *k*-mers serve as keys and counts and sample IDs are associated data. Initially, the heap stores the lexicographically lowest *k*-mers from every sample. Then, a *k*-mer from the top of the heap is extracted, followed by removing the same *k*-mers originating from the other samples. The heap is amended by inserting lexicographically next *k*-mers from the proper samples.

Before adding a new row to *M*, two filtering criteria are verified. First, if *F* is provided, the *k*-mer must be present in *F*. Since *F* is represented as a compatible KMCBD file, the distribution of its *k*-mers into bins is the same. Such verification requires a single linear traversal over *F*’s *k*-mers. Second, if *η* is the number of samples for which *c*(*x, j*) ≥ *τ*, we require that 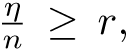 where *τ* (1 by default) and *r* (0 by default) are user-provided parameters.

The matrix *M* is stored in KMCDB, and optionally, in a text file. Together with the KMCDB file, we also keep additional data to simplify later normalization, such as the sums of the retained counts for each sample.

Merging is performed by *t* threads. Each thread receives a unique bin ID from a pool and processes *k*-mers present in the bin with this ID from all the samples. When there are no bins to process, the thread ends. To balance the computation load, the bins are sorted in descending order by the number of *k*-mers they contain before starting work.

### 2.4 Analyzing the matrix

The final stage involves analyzing the matrix *M*. The user can select one or more scenarios here. In most cases, the input to the following methods can be a matrix *M* with raw counts or the matrix normalized *M* ^norm^. Currently, we support three normalization algorithms: DESeq2, frequency count, and quantile. The normalized matrix can be optionally output to a text file.

When calculating correlation (Kendall Tau, Pearson, Spearman algorithms), performing differential *k*-mers analysis (ANOVA, DIDS, Signal to Noise ratio, *t*-test, Wilcoxon-rank sum), or determining the entropy, the *k*-mers can be processed independently, so we need just a single row of *M*, and possibly a phenotype, at a time. Therefore, each bin is traversed by extracting consecutive rows from the matrix with the *k*-mers and their corresponding counts, and then doing the computations. The results (e.g., correlation values) are stored in a proper output file. If *p*-value correction is required, the values are temporarily kept in vectors (in main memory) and corrected (we support Bonferroni, Benjamini-Hochberg, Benjamini-Yekutieli, and Holm-Bonferroni algorithms) in a second pass.

A parallel processing employs a similar bin-oriented scheme as merging, i.e., each thread processes a single bin of matrix rows. Parallel data storage in KMCDB utilizes the properties of KMCDB streams (see Section 2.5). Storing results in text files utilizes thread-local buffers, which are filled with consecutive results and periodically flushed to disk exclusively. Thus, the order of *k*-mers in text files is neither preserved nor specified.

Since the correction is a sequential task, one thread is responsible for the full computation of corrections for the entire single differential *k*-mers analysis method (e.g., *t*-test). Corrected *p*-values are then transferred from memory to a disk with *t* threads in a similar parallel manner as in the previous stages.

For dimensionality reduction (we support PCA and UMAP methods), we need to keep the whole *M* ^norm^ matrix in main memory. This can result in large memory requirements. In our implementation, we employed third-party programming libraries for both PCA and UMAP.

### 2.5 KMCDB

KMCDB is a file format designed for storing *k*-mers with associated data across multiple samples. From the MKMC perspective, it may be viewed as a matrix with *k*-mers as rows, samples as columns, and counts as entries. In addition to integer values, KMCDB also supports real numbers or tuples (a feature not utilized by MKMC).

We designed KMCDB with two primary objectives:

1. **Multithreading support** – aiming for fast read/write operations in a streaming fashion by multiple threads.
2. **Flexible data representation** – supporting multiple ways of storing data and allowing new representations to be added in the future.

To achieve this, access to KMCDB is split into two layers. The high-level layer manages data representation, versioning, and metadata, while the low-level, *archive* layer is a custom container-based database responsible for disk I/O and synchronization. The details of the archive are described in the next section. Conceptually, the archive is composed of *streams* of data, each consisting of *parts*. The archive can be viewed as a single directory with subdirectories (archive streams), containing numbered files (archive parts) (Fig. 2, middle part). A concatenation of these files (in the order defined by their numbers) forms a stream. The part is a transaction unit between the high-level and low-level layers.

**Fig. 2:**
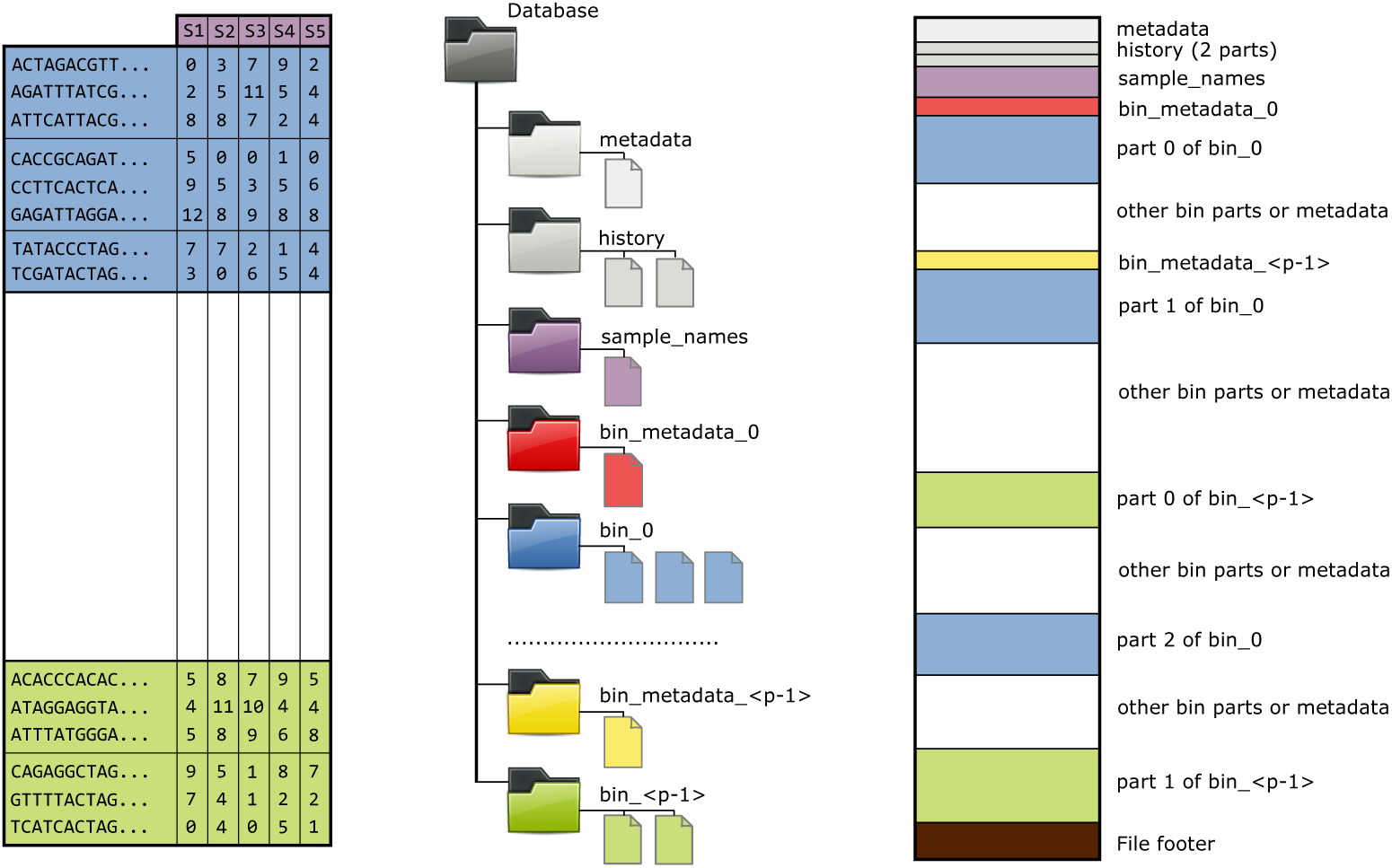
KMCDB file design. Illustration of how the MKMC matrix is represented in KMCDB format using a container-based, *archive*, database. The left part shows a matrix of counts related to *k*-mers for five samples (S1,…, S5). The *k*-mers are organized in bins (marked by the same background color), and within each bin, they are sorted. Each bin is divided into several parts. The split is made when a buffer collecting *k*-mers becomes full. The middle part illustrates the logical organization of the KMCDB database, which is composed of data in streams, similar to a directory structure. The example shows one of the ways *k*-mers can be stored, i.e., without splitting into prefix and suffix. The right part shows how this is organized in a single file in *archive* format.

Accesses to KMCDB are buffered, and the buffer stores many rows. In writing mode, if the buffer is full, it is sent to the archive as a single part. Symmetrically, in a reading mode, when all rows in the buffer are read, the next part is received from the archive. Each archive stream may be processed independently, which is the basis of multithreaded processing in MKMC. Specifically, the matrix rows are partitioned into multiple bins based on *k*-mer signatures. This splitting is performed during the *k*-mer counting stage. All *m*-mers (for *m < k*) are extracted from a *k*-mer, their hash values are computed, and the minimum is selected as a signature of a *k*-mer. The bin number is computed as the signature modulo the number of bins.

The organization of KMCDB data is described in the *metadata* stream, which defines: KMCDB version, *k*-mer length, number of columns (samples), number of bins, number of bytes used to represent a single matrix entry (in the case of tuples as entries, it is a list of values), signature info (length, selection scheme, bin mapping), and representation of the data. Two KMCDB files are *compatible* if they have the same *k*-mer length, signature info, and the number of bins. Signature selection scheme describes how, for a given *k*-mer, its signature is computed, while bin mapping defines how the bin number is computed for a signature. Currently, KMCDB supports a single signature selection scheme, single signature bin mapping, and two methods of representing the data. In the first one, *k*-mers are stored in a 2-bits per symbol (mapped as A→0, C→1, G→2, T→3), followed by the row entries. In such a case, there is a separate stream in the archive for each bin. The second one is similar, but *k*-mers are split into prefix and suffix, just as is done in KMC 3. In such a case, instead of a single stream for bin data, there are two streams. One contains the data required to reconstruct *k*-mer prefixes, and the second contains *k*-mer suffixes followed by the row entries. In both cases, there are additional metadata streams for each bin. Currently, it contains only the number of *k*-mers in the given bin.

The history stream records auxiliary information, such as the command line used by the tool that created the given KMCDB, memory usage, the time between KMCDB open and close, and the run environment (including hardware and software information). All this data is stored as a single part in the history stream, forming a single history item. If a new file in KMCDB format is constructed based on another KMCDB, all previous history items are rewritten to the new file. Names of samples/columns are stored in a separate stream. If they are not provided, the stream is absent.

In MKMC, KMCDB is opened in a streaming mode, i.e., only a part of the data is actually loaded into memory. Nevertheless, KMCDB can also be opened in a query mode that enables random access. In the future, we plan to extend KMCDB by adding new representation modes for both *k*-mers and associated values.

### 2.6 Archive

The container-based database, *archive*, was first developed for the VCFShark tool [14] and has since been used in various projects, undergoing significant extensions. It allows for the registration of many streams, and each stream can be composed of many parts. The user can add parts (optionally accompanied by a single 64-bit integer, part-level metadata) of the streams in any order from multiple threads.

Technically, the database is a single file composed of parts of streams in the order they are added to the database. A footer concludes the file. For each stream, it stores its name, number of parts, offsets to each part in the file, part size, and the stream-level 64-bit metadata field. At the very end, the archive stores a few internal parameters, such as the format version and the method used to store integers. The right part of Fig. 2 illustrates how the KMCDB streams (middle part of the figure) could be placed in the file. A brown box represents the file footer.

While reading, the user can access any part of any stream in any order. One can also read the data from each stream sequentially, part by part, or in smaller chunks.

The archive database allows for registering streams at any time, so the database structure does not need to be known initially. The streams can store any binary data. Moreover, the database can be updated by adding new streams or new parts of the existing streams. This allows extending the applications using the database without breaking backward compatibility.

## 3 Results

### 3.1 MKMC Computational Performance

All experiments were made on a server equipped with AMD Epyc 9654 CPU (96 cores, base clock 2.4 GHz), 2474 GB RAM and 16 TB NVME disk. We evaluated MKMC against Kamrat [7] and kmtricks [15] on two RNA-seq datasets: GSE216369 (16 samples; 224 GB) [16] and GSE132040 (933 samples; 12.5 TB) [17] (see Supplementary Section 13.4 for additional information). Unless stated otherwise, multi-threaded tools were run with 32 threads.

The main results are presented in Figure 3. The first step of the analysis is always to build a matrix of *k*-mer counts for each sample (see Fig. 3ab). MKMC accomplished this in 118 s (GSE216369) and 7,164 s (GSE132040). kmtricks was remarkably slower: 321 s and 19,080 s, respectively. Kamrat is not equipped with the matrix building step, so, following the authors’ suggestions from GitHub repository, we used Jellyfish [18] and joinCounts from the DE-kupl pipeline [19] for that, and Kamrat itself to index the matrix. This took 7,805 s for the smaller dataset and did not complete in 24 h for the larger one; these were the only cases when matrices were built in a text, not binary format. Memory consumption of all the algorithms was similar. For the smaller dataset, MKMC won (15.7 GB vs. 24.1 GB and 16.4 GB for kmtricks and Kamrat), while for the bigger one kmtricks was more frugal 12.6 GB vs. 17.0 GB).

**Fig. 3:**
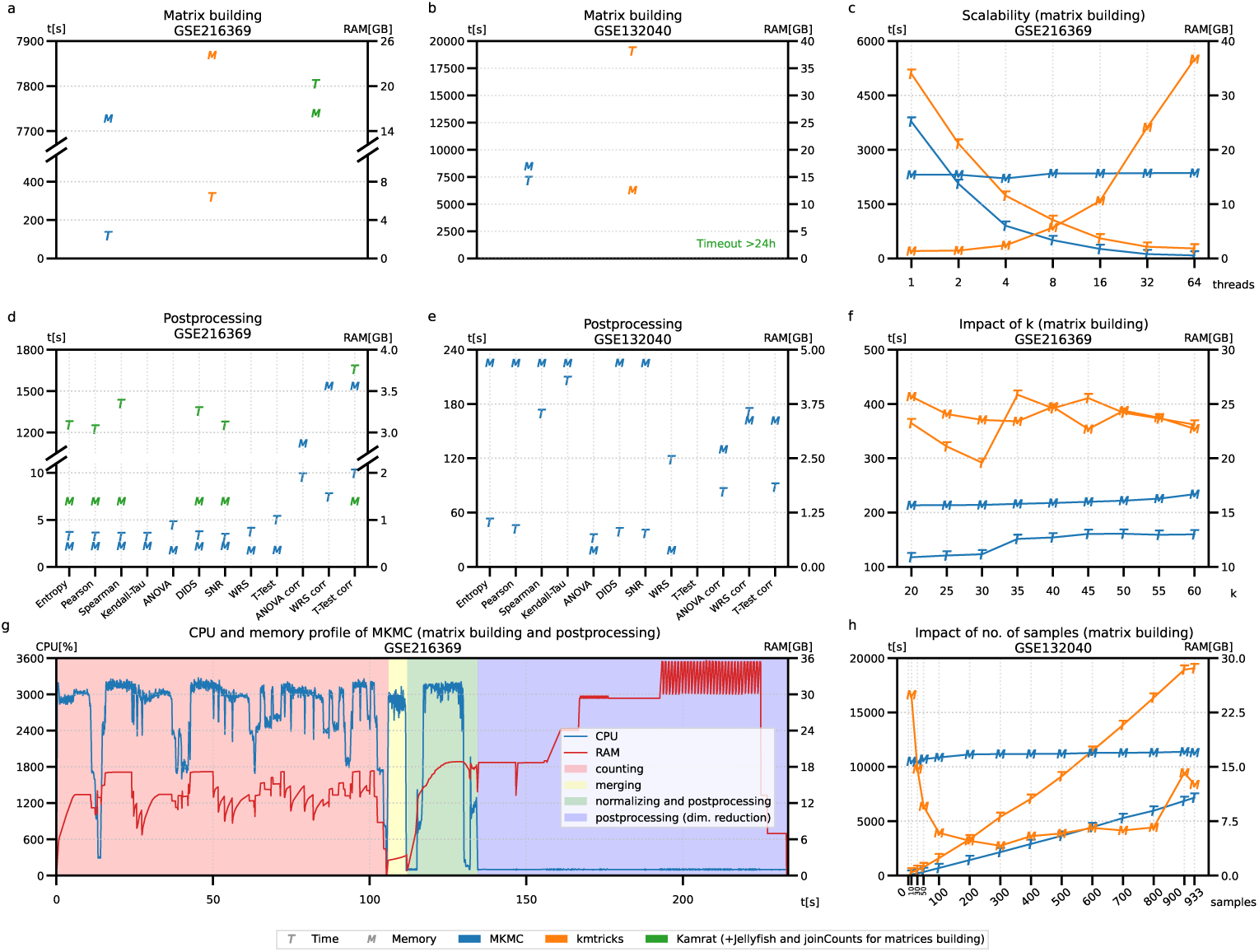
**MKMC is in general faster and requires less memory than competitors.**Time and memory performance of MKMC and the competing tools for datasets GSE216369 (16 samples; 224 GB), GSE132040 (933 samples; 12.5 TB). a, b — Initial building of *k*-mers (*k* = 27) counts matrices. For GSE132040, we were unable to count k-mers with Jellyfish in 24h to start actual building with Jellyfish, joinCounts, and Kamrat pipeline. c — For MKMC and kmtricks, the building was performed for various numbers of threads. d, e — MKMC was used to postprocess the previously built binary matrix and obtain various results; if necessary, counts were normalized with the frequency count method. f — The building was performed for various values of *k*. g — MKMC was utilized to perform all the tasks in a single run: building, normalizing, and postprocessing. The plot shows CPU utilization (32 threads) and memory usage. h — The building was performed on a different number of samples randomly chosen from GSE132040; kmtricks required to increase ulimit.

Once the matrix is ready, various analyses can be performed. Fig. 3de shows the time requirements and memory use of MKMC and Kamrat for GSE216369 and MKMC for GSE132040. One can see that most MKMC analyses are completed within a few seconds for the smaller dataset and within a few minutes for the larger dataset, while Kamrat requires above 20 minutes. The memory needed for both the tools was always less than 5 GB. In most cases, MKMC processed *k*-mers in parallel, in a streaming fashion, which explains its low time and memory requirements. When *p*-value correction was required, all *p* values had to be loaded into memory, and this part of the processing was sequential, which explains both the longer running times and the higher memory use.

In the following experiments, we focused on the scalability of matrix construction and the impact of the *k*-mer size in MKMC and kmtricks. Fig. 3c shows that the running times drop rapidly as the number of threads increases. The situation with memory use, however, is different. For MKMC, it is constant, which is caused by the internal use of KMC with a fixed upper memory bound. kmtricks’ memory usage depends strongly on the number of threads.

The impact of *k*-mer length is moderate in both tools (Fig. 3f). The only interesting observation is the growth of running time when crossing the *k* = 32 boundary. This relates to how *k*-mers are represented internally. For *k* ≤ 32, they are stored in a single 64-bit word, and for 32 *< k* ≤ 64, they require two such words, which slows all operations on them.

The last experiment in this series focused on the influence of the number of input samples. (Fig. 3h). As it grew, both MKMC and kmtricks scaled almost linearly in running time. The memory use remained almost constant for MKMC across the full range of input sizes, but for kmtricks it grew rapidly when exceeding 800 samples, and for less than 100 samples.

To provide a deeper insight into MKMC, we obtained a thorough profile of MKMC steps, including snapshots of memory and CPU utilization (Fig. 3g). The CPU utilization strongly depends on the processing stage. In the counting, when KMC is employed, it is around 3000%, which is almost perfect, given that 32 threads are employed. Then, in merging, it is similar, and in analyzing (postprocessing) again exceeds 3000%. Finally, in the dimensionality reduction step, it is 100%. MKMC uses 3rd-party libraries for PCA and UMAP. Their processing is serial, which explains the phenomenon. Regarding memory consumption, the dimensionality reduction dominates the whole pipeline. Once again, this is the effect of the external programming libraries used here.

Another group of experiments aimed at both verifying resource consumption and comparing the quality of MKMC results with that of mapping-based gene-level experiments. Figure 4 shows results of experiments that will be described in detail in Sections 3.2 and 3.3, respectively. They include full pipelines, from trimmed reads to results generation. As both gene-level pipelines differ in the last step, which is running a Python script, they yields similar results. MKMC can generate results in less than 3 minutes, compared with more than 1 hour for the mapping-based approach. It also includes fewer steps and no need to install Python packages. Memory requirements are similar in both pipelines.

**Fig. 4:**
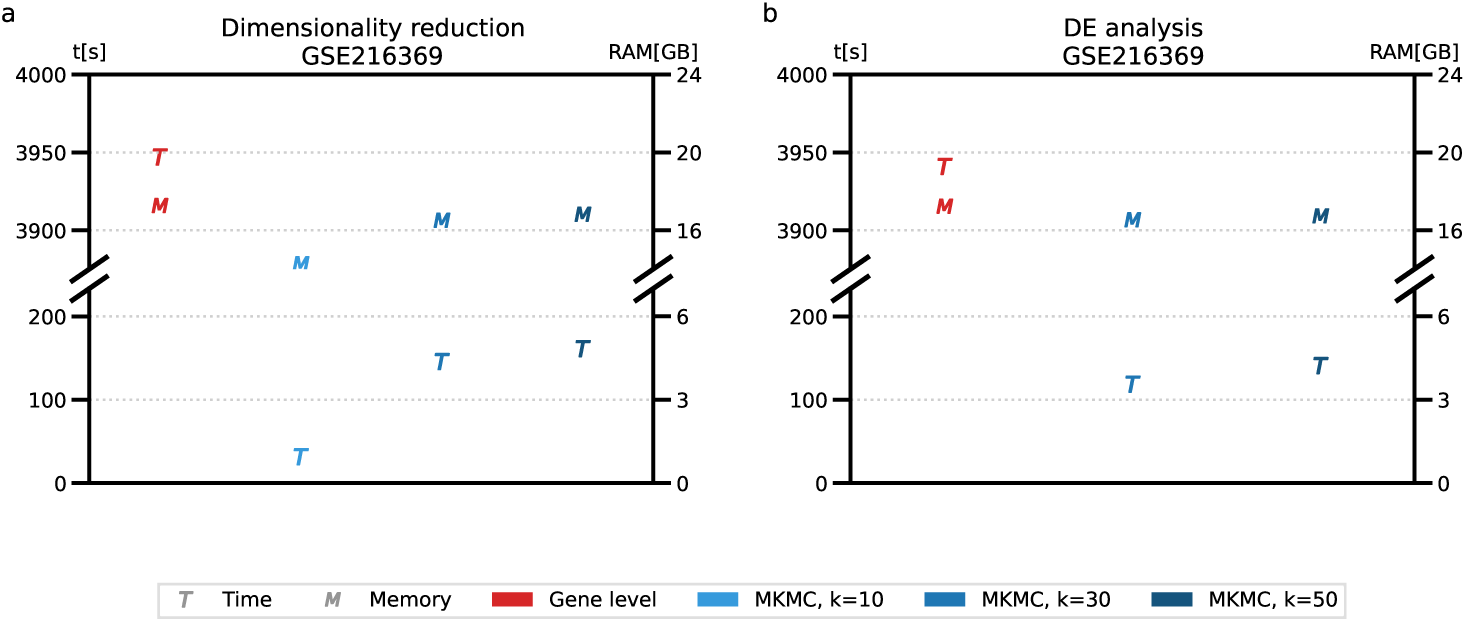
MKMC is much faster and requires less memory than mapping-based approach. Comparison of time and memory performance for MKMC and gene-level (mapping-based) analysis for a dataset GSE216369. The plots show results of: a — dimensionality reduction and b — differential expression analysis.

### 3.2 PCA and UMAP of K-mers Reveal Sex Dimorphism Comparable to Traditional Alignment

To assess the impact of the alignment-first versus alignment-last approaches for unsupervised visualization of biological variation, we compared two pipelines: (1) a traditional alignment-based RNA-seq analysis pipeline using STAR, and (2) a novel k-mer-based pipeline using MKMC. Both pipelines were applied to bulk RNA sequencing data from the African Turquoise Killifish liver tissue[16]. This dataset consists of 16 samples, 8 samples from males and 8 samples from females. Half of the animals were fed using ad libitum diet and the other half were fed using a dietary restriction diet. For our analysis that compares between males and females, we pooled the animals in both dietary regimens. Before performing unsupervised visualization techniques, including Principal Component Analysis (PCA) and Uniform Manifold Approximation and Projection (UMAP), the count matrices from both methods were frequency-normalized[20, 21]. This normalization step helped ensure comparability of expression values between the two approaches by adjusting for sequencing depth differences and accounting for sample-to-sample variability in library size, preventing larger or smaller sequencing libraries from dominating purely due to depth. Here we investigated whether UMAP and PCA could effectively separate male and female samples in both pipelines, providing insights into how biological variation is captured by each approach as seen in Fig. 5.

**Fig. 5:**
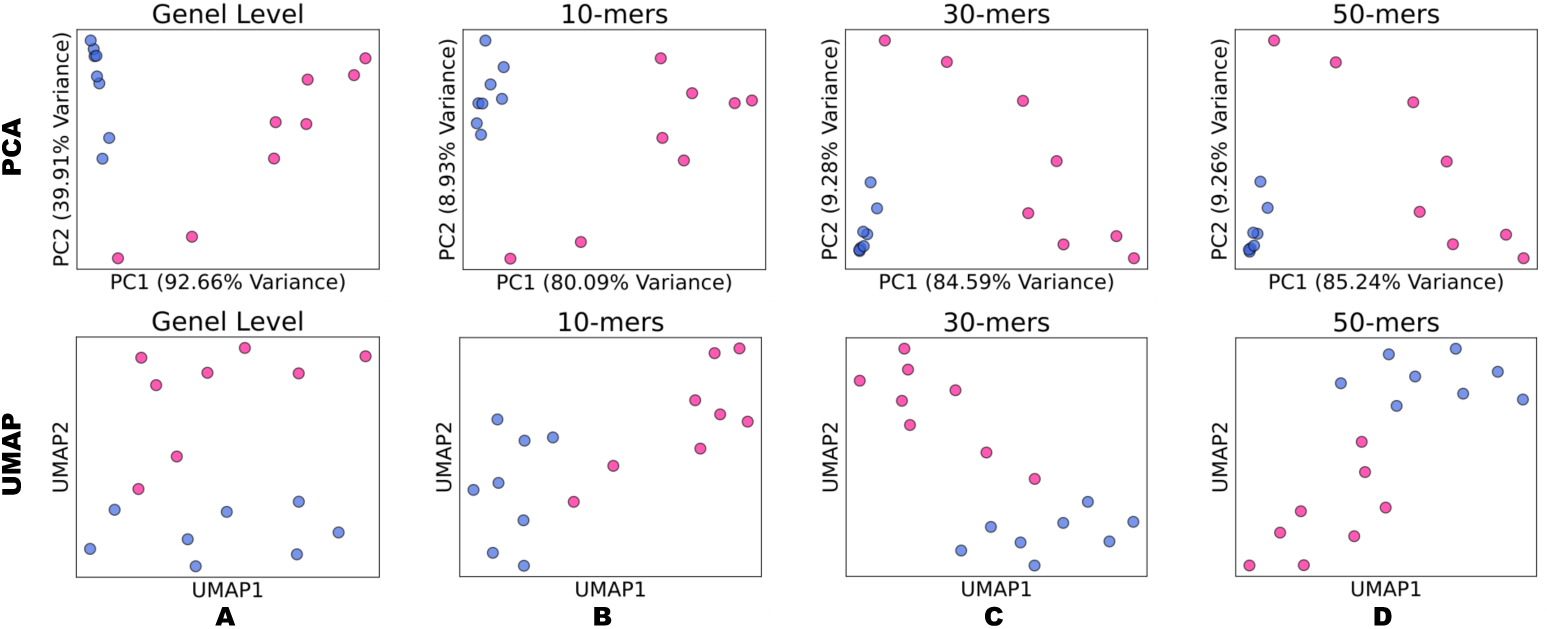
PCA and UMAP Reveal Biological Variation in Gene-Level and K-mer Analyses. PCA and UMAP were used to compare biological variation across gene-level features (traditional RNA-seq) and k-mer-based features (10-mer to 50-mer) generated using MKMC. PCA consistently showed PC1 and PC2 explained the majority of variance, while UMAP revealed similar separation patterns. Higher k-mer sizes resulted in fewer features, with minimal impact on variance explained. The blue color represents male samples and the pink samples represent female samples.

For the gene-level analysis, which consisted of 25,122 features, PCA revealed that the first two principal components (PC1 and PC2) captured 96.65% of the total variance, effectively separating biological conditions with minimal technical variation. UMAP demonstrated similar separation patterns, further confirming this separation.

In k-mer-based analyses, increasing k-mer size reduced the number of features but did not substantially affect the variance explained by PC1 and PC2. For 10-mers (524,800 features), PCA captured 89.02% of the total variance, while 20-mers (18,276,797 features) explained 91.68%. At 30-mer (16,478,913 features), the variance captured increased to 93.87%, and at 40-mer (14,890,416 features) and 50-mer (13,337,478 features), the variance reached 94.42% and 94.5%, respectively. UMAP visualizations consistently reflected clear separation of biological conditions across all k-mer sizes, mirroring the trends observed in PCA. These results highlight the robustness of k-mer-based approaches for identifying biological variation, even with fewer features at higher k-mer sizes.

### 3.3 Direct K-mer Analysis Recapitulates Gene-Level RNA-Seq DE Patterns

Furthermore, we evaluated the performance of the traditional alignment-based method and the k-mer based approach through differential expression (DE) analysis on bulk RNA sequencing data from the same liver tissue used for the dimensionality reduction. We compared the traditional DE method (based on sequence alignment using STAR) with MKMC that employs k-mers of size either 30 or 50, to reveal the similarities and differences in the DE gene identification and the impact of the k-mer length on the results.

Using a traditional pipeline we aligned the RNA-seq reads to a reference genome by using STAR aligner and then used the featureCounts package to generate a count for each gene[10, 22]. Finally, methods such as DESeq2 and Limma-Voom are used to identify DE genes[23, 24]. This method can further allow for the detection and differential abundance quantification of indels and splice junctions. However, we performed differential expression analysis using the Wilcoxon rank-sum test following frequency count normalization. The Wilcoxon test is a non-parametric method that does not assume a specific distributional form, making it well-suited for normalized k-mer count data, which may deviate from the negative binomial assumptions underlying traditional RNA-seq methods. Additionally, prior studies have shown that rank-based methods can provide robust control of false discovery rates and competitive performance in large-sample settings, particularly when distributional assumptions are violated[25]. The Wilcoxon rank-sum test identified 1,663 DE genes, leveraging its robustness for FDR control in high-dimensional datasets, unlike methods like DESeq2 or Limma-Voom that rely on parametric assumptions[25]. In contrast, the k-mer-based method directly detects DE k-mers from raw reads without the need for alignment to a reference genome.

Using the same killifish liver tissue dataset as in our unsupervised analysis, we compared differentially expressed genes (DEGs) identified by the traditional alignment-based method with those found using our k-mer–based approach. As shown in Fig. 6, we evaluated the statistical significance of overlap in DEGs using the hypergeometric test. In the control (“Permuted”) condition, we randomly shuffled gene labels to test whether any observed overlap was due to chance. In contrast, the“Ranked” condition uses the actual ranked list of genes obtained from the Wilcoxon rank-sum test on k-mer counts. The ranked k-mer results (both 30-mers and 50-mers) show significant overlap with the DEGs identified by the alignment-based method, while the permuted control does not, indicating that the k-mer–based approach captures meaningful biological signal rather than random noise as seen by Fig. 6.

**Fig. 6:**
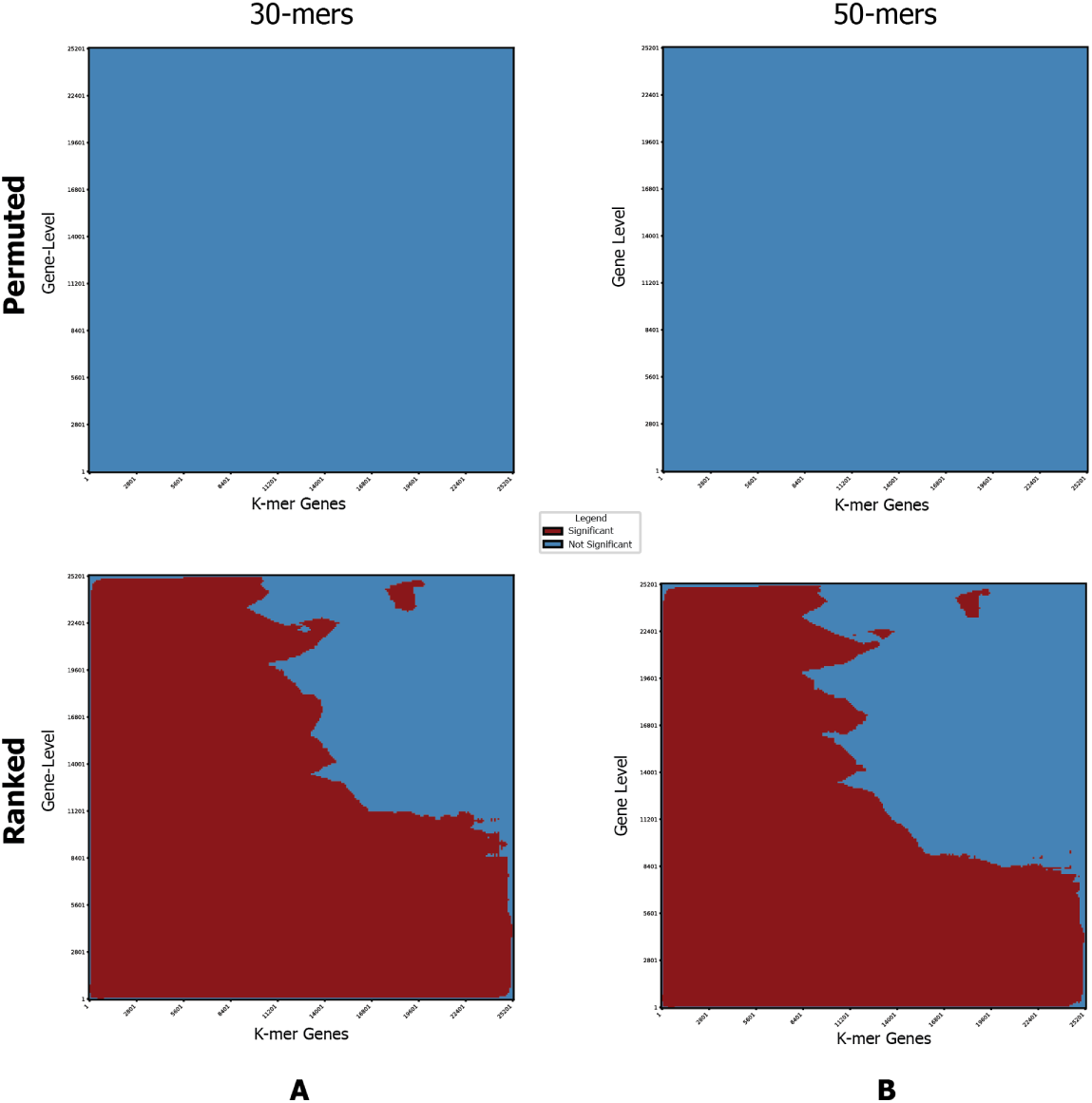
High Concordance Between Traditional Alignment and K-mer Based Methods in DE Analysis. Comparison of differential expression (DE) analysis between the traditional gene-level alignment method and k-mer-based methods (30-mers and 50-mers) using the Mann-Whitney U (also known as the Wilcoxon rank-sum) test. **(A)** Hypergeometric test results showing overlap between DE genes identified by the traditional alignment-based method and 30-mer k-mer–based methods. “Permuted” refers to a control where gene labels are randomized, while “Ranked” refers to actual DE gene rankings from Wilcoxon analysis. Only the Ranked comparisons show statistically significant overlap. **(B)** Same as (A), but using 50-mers. In both cases, k-mer–based DEGs show high concordance with alignment-based DEGs, confirming the biological relevance of the k-mer–based method. Both sets of k-mer lengths identify comparable sets of DE genes as with the traditional alignment method.

**Fig. 7:**
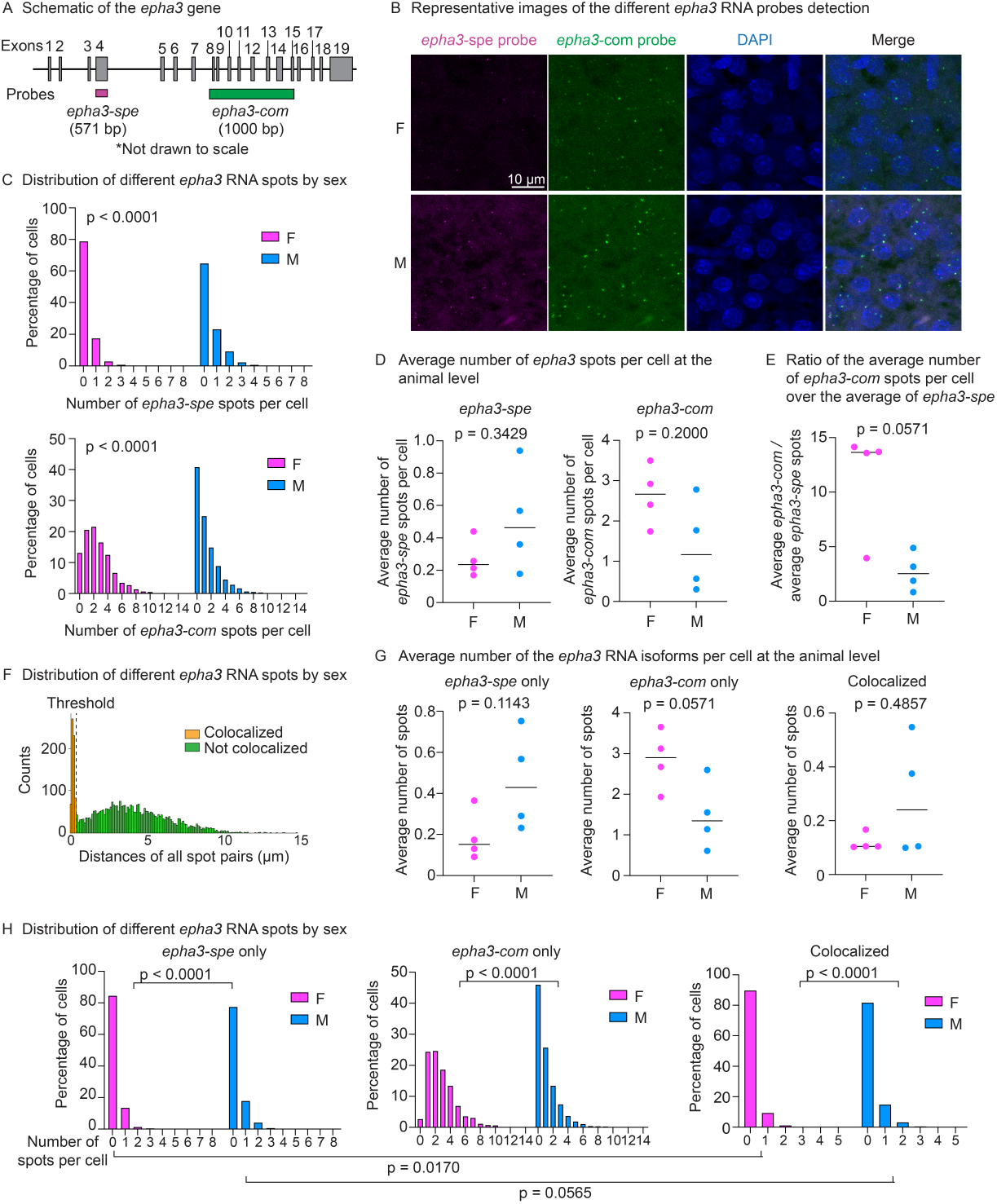
Sex-Specific Expression Patterns of *epha3* Validated via RNA In Situ Hybridization. A) Schematic of the *epha3* gene, which has 19 exons (gray boxes). The male-enriched RNA sequence includes 571 base pairs (bp) aligned to exon 4. The ‘*epha3-spe*’ probe anneals to this 571-bp region (purple). The RNA sequence detected in both males and females align to several exons, including exon 8-15, and the ‘*epha3-com*’ probe anneals to a 1000-bp region spanning from exon 8 to exon 15. This size of the exons and introns (lines connecting the exons) are not drawn to scale. B) Representative images of the different *epha3* RNA probes detection. F, female; M, male. Scale bar, 10 µm. C) Distribution of the number of *epha3-spe* spots per cell (top graph) and the number of *epha3-com* spots per cell (bottom graph), separated by sex. Each distribution was pooled from 4 animals of each sex (two fields of view was imaged for each animal), totaling 2059 cells for females and 2001 cells for males. Statistics was performed using Kolmogorov-Smirnov test, comparing the male and female distributions. D) The average number of *epha3-spe* (left) or *epha3-com* (right) spots per cell at the animal level (n = 4 for female, n = 4 for male). For each animal, the average number of either *epha3* RNA spot type was calculated by dividing the total of *epha3-spe* spots or *epha3-com* spots by the total number of cells, respectively. Statistics was performed using Mann-Whitney test. E) The ratio of the average number of *epha3-com* spots per cell over the average number of *epha3-spe* spots per cell for each animal. Statistics was performed as in panel D. F) The distribution of all distance pairs between each ‘*epha3-spe*’ spot and ‘*epha3-com*’ spot in each cell. Distances are binned every 0.075 µm. Dotted line denotes the distance threshold (0.375 µm). Spot pairs with distances shorter than this threshold were classified as ‘colocalized’ (orange) and those farther than this threshold (green), classified as ‘non-colocalized’ (i.e., ‘*epha3-spe*’-only or ‘*epha3-com*’-only). G) The average number of ‘*epha3-spe*’-only (left), ‘*epha3-com*’-only (middle), and colocalized spots (right) per cell at the animal level. Statistics was performed as in panel D. H) Distribution of the number of ‘*epha3-spe*’-only (left), ‘*epha3-com*’-only (middle), and colocalized spots (right) per cell separated by sex. Each distribution was pooled from 4 animals of each sex as in panel C. Statistics was performed using Kolmogorov-Smirnov test, either comparing the male and female distributions or comparing the *epha3-spe*’-only and the colocalized spots distributions for males or females.

To quantitatively assess the concordance between k-mer–based and traditional alignment-based differential expression analyses, we constructed a contingency table (Table 1; results for k-mer-based method may insignificantly vary between runnings due to BWA indeterminism) comparing gene-level significance calls between the two methods. The table reveals a strong overlap: 1,474 genes were identified as significantly differentially expressed by both approaches. In contrast, only 189 genes were found to be significant exclusively by the alignment-based method, and 2,472 genes exclusively by the 30-mer method. The vast majority of genes (20,987) were classified as not significant by either method. This asymmetric distribution—with a disproportionately large intersection in the significant/significant cell—strongly suggests that the k-mer–based approach reliably recovers biologically relevant signals captured by alignment-based analysis, while also potentially uncovering novel signals missed by alignment-dependent pipelines.

**Table 1:** Contingency table comparing DE results from 30-mers and gene-level analyses.

|  |  | Gene Space |  |
| --- | --- | --- | --- |
|  |  | Significant | Not Significant |
| K-mer Space | Significant | 1,474 | 2,472 |
|  | Not Significant | 189 | 20,987 |

Among the 2,472 genes identified as significant by MKMC—defined as genes with at least one significant 30-mer alignment—were *Nfatc2ip* (254 significant 30-mers), *Epha3* (813 significant 30-mers), and *LOC107389788* (691 significant 30-mers). These examples illustrate the power of our k-mer–based approach to detect diverse transcriptomic events, including alternative 3^′^ UTR usage (*Nfatc2ip*), alternative transcription start sites (*Epha3*), and potential alternative splicing (*LOC107389788*).

### 3.4 Validation of MKMC Predictions Using RNA In Situ Hybridization

To validate the findings of MKMC, we used a sensitive RNA *in situ* hybridization method called Hybridization Chain Reaction (HCR) for detecting specific mRNA sequences in male and female killifish liver sections. As an example, we designed two sets of HCR probes for the *epha3* gene, one against the male-enriched sequence detected by MKMC (‘*epha3-spe*’) and another against the sequences detected in both sexes (‘*epha3-com*’) (Figure 4A). We observed that male livers had more cells expressing a higher level of male-enriched RNA sequence than female livers (Figure 4B-C, ‘*epha3-spe*,’ p *<* 0.0001, Kolmogorov-Smirnov test). In contrast, the common mRNA sequence is more highly expressed in female livers than male livers (Figure 4B-C, ‘epha3-com,’ p *<* 0.0001, Kolmogorov-Smirnov test). Consistent with our predictions by MKMC (Figure 3), the ratio of the two mRNA sequences differed at the animal levels (p = 0.0571, Mann-Whitney test), with a stronger expression preference on the common sequence in female livers than male livers (Figure 4D-E).

Next, we performed colocalization analysis to study the *epha3* transcript isoforms. A colocalized spot would correspond to an RNA molecule that contained both the male-specific and the common sequences. A non-colocalized spot could suggest a shorter mRNA isoform with only part of the sequence in the longer isoform (though 8% of non-colocalized spots could be due to missed detection)[26]. A ‘*epha3-spe*’-only RNA would contain a more 5^′^-end sequence of the longer isoform, whereas a ‘*epha3-com*’-only RNA would have a more 3^′^-end sequence. To classify the colocalized spots, we identified a distance threshold based on the separation in the distribution of all distance pairs between each ‘*epha3-spe*’ spot and ‘*epha3-com*’ spot in each cell (Figure 4F, dotted line) and denoted any spot pair with a distance less than this threshold as ‘colocalized’ (Figure 4F, orange bins). Of all the cells that expressed any *epha3* RNA (i.e., excluding cells with no detected *epha3* HCR spots), the male livers expressed a higher average of the ‘*epha3-spe*’-only RNAs per cell (a more 5^′^-end sequence) than female livers (Figure 4G, p = 0.1143 by Mann-Whitney test at the animal level, and Figure 4H, p *<* 0.0001 by Kolmogorov-Smirnov test at the distribution level), whereas female livers expressed higher average of the ‘*epha3-com*’-only RNAs per cell (a more 3^′^-end sequence) than male livers (Figure 4G, p = 0.0571 by Mann-Whitney test at the animal level, and Figure 4H, p *<* 0.0001 by Kolmogorov-Smirnov test at the distribution level). In addition, the male livers had relatively similar expression distributions of the ‘*epha3-spe*’-only isoform and the longer isoform (‘colocalized’ spots) (Figure 4H, p = 0.0565, Kolmogorov-Smirnov test), whereas the female livers had a slightly lower level of the longer isoform than the ‘*epha3-spe*’-only isoform (Figure 4H, p = 0.0170, Kolmogorov-Smirnov test). Together, our analysis suggested that male and female livers expressed different epha3 isoforms at different levels, with higher expression of a more 5^′^-end RNA isoform in males and higher expression of a more 3^′^-end RNA isoform in females.

### 3.5 Transcriptomic Age Prediction

Accurate biological age prediction is critical for understanding aging and its associated biological mechanisms. Traditional gene-level methods for age prediction often rely on RNA-seq-based expression data, where the relative abundance of annotated genes serves as the foundation for building predictive models. Here, we introduce a k-mer-based approach that circumvents the limitations of relying on annotated genomes and directly utilizes sequence-level information to predict transcriptomic age.

For our age prediction analysis, we used male retina samples from the African Turquoise Killifish collected at four age points: 1.5, 3.0, 4.5, and 6.0 months, with three biological replicates per time point. As shown in Fig. 8, the k-mer–based method achieved predictive performance comparable to traditional gene-level model using Leave One Out Cross Validation (LOOCV). Using Elastic Net regression, we trained age clocks on both k-mer counts and gene expression data. The k-mer–based model achieved a Pearson correlation of 0.900 and a mean absolute error (MAE) of 0.607 months between predicted and chronological age—closely matching the gene-level model’s correlation of 0.887 and MAE of 0.695 months. These results underscore the effectiveness of k-mers as predictive features, capturing both coding and non-coding transcriptomic signals relevant to aging.

**Fig. 8:**
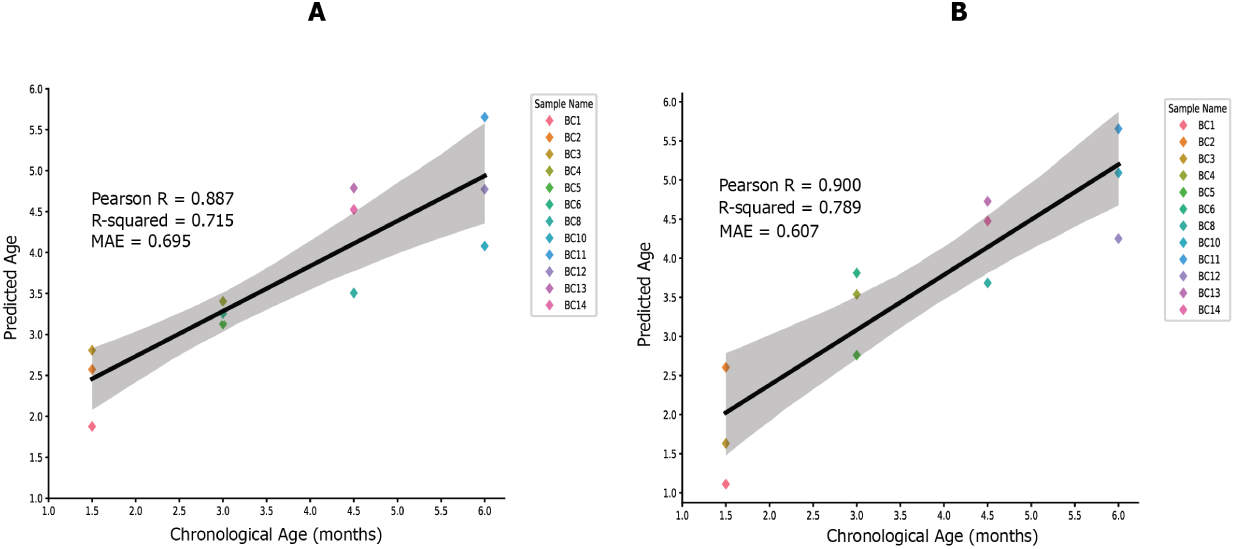
K-mer based Age Prediction Matches Performance of Gene-Level Age Prediction. Comparison of age prediction models using (A) gene-level and (B) k-mer-based (30-mers) approaches. The scatter plot illustrates the correlation between predicted and chronological ages across both methods, demonstrating the comparable performance of gene-level age prediction and k-mer-based age prediction. Both approaches utilized Elastic Net regression, highlighting the robustness of k-mer features as an alternative to traditional gene-level features. While the feature space at the gene level is about 25,000, the 30-mers analysis comprised of 696,014 features. These were canonical 30-mers with at least 500 counts in all the samples.

**Fig. 9:**
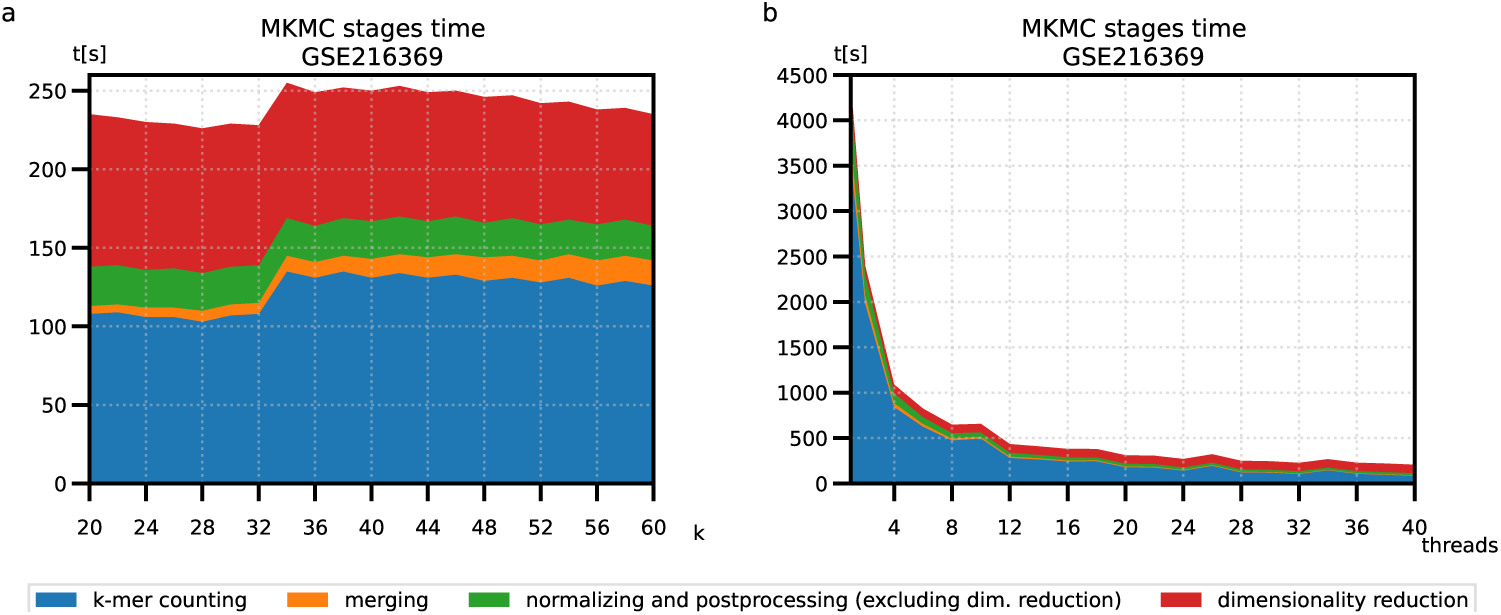
Time of various MKMC stages depending on k-mer length and number of threads. MKMC was utilized to perform all the tasks in a single run: building the matrix and filtering, normalizing, and the matrix postprocessing. The plot shows time of its main stages (k-mers counting, merging — to build a matrix and filter) depending on a value of of *k*, and the number of threads.

By directly comparing k-mer-based and gene-based models performance, we demonstrate the flexibility and broad applicability of k-mer features for biological age prediction. This approach aligns with our overarching goal of expanding transcriptomics beyond traditional alignment-based workflows, paving the way for novel insights into aging biology. We also implemented a cross-validation parameter in MKMC to make machine learning applications using k-mer more accessible to the community.

## 4 Discussion

K-mer–based approaches represent a general and powerful framework for sequence analysis, enabling the decomposition of biological sequences into fixed-length substrings that can be efficiently counted and compared without reliance on reference genomes [1, 4]. This paradigm has been widely applied across genomics and transcriptomics, offering advantages in computational efficiency, scalability, and the ability to capture both known and novel sequence features [1]. RNA sequencing (RNA-seq) represents a prominent application domain in which the advantages of k-mer–based methods are particularly evident. Traditional RNA-seq analysis tools have enabled major advances in transcriptomics but face important limitations in flexibility, interpretability, and scalability [9, 27]. The comparative summary in Table 2 highlights these trade-offs across popular alignment-based methods such as STAR, RSEM, Salmon, and DESeq2 [10–12, 23, 28, 29]. These pipelines are built on the assumption of a complete, high-quality reference genome, making them less effective for non-model organisms or poorly annotated systems [5, 6]. They also rely on alignment heuristics—such as collinearity and optimal gap scoring—that can influence transcript abundance estimation and obscure biologically meaningful variation, including alternative splicing and isoform-specific regulation [2–4]. To address these limitations, k-mer–based methods have emerged as flexible alternatives for transcriptomic analysis. As summarized in Table 3, these approaches have been successfully applied to tasks ranging from alternative splicing detection (e.g., Kissplice) to more recent scalable, multi-condition analyses such as sc-SPLASH [7, 13, 15, 30–41]. However, existing tools often involve trade-offs in speed, usability, or analytical scope, limiting their broader adoption. In this work, we build on these advances by introducing MKMC, a centralized and scalable toolkit designed to leverage the strengths of k-mer–based analysis while addressing the limitations of existing approaches. While MKMC can operate in a fully reference-free mode, the biological analyses presented here incorporate reference-based filtering and annotation to facilitate interpretation of k-mer signals. As shown in Table 4, MKMC integrates multiple analytic modules—including k-mer counting, normalization, differential expression testing, and dimensionality reduction—into a single, user-friendly framework that improves computational efficiency and analytical breadth compared to prior tools. This design enables comprehensive analysis of sequencing data without requiring alignment as a primary processing step, while still allowing downstream integration with reference-based annotations when needed. To benchmark MKMC’s utility, we first applied unsupervised visualization techniques such as PCA and UMAP to k-mer frequency profiles. These analyses recapitulated known biological distinctions, such as sex differences in liver transcriptomes of the African turquoise killifish, with patterns comparable to gene-based analyses derived from alignment pipelines. This consistency was observed across different k-mer sizes, indicating that k-mer representations preserve biologically meaningful structure despite differences in feature dimensionality. We next performed differential expression analysis using a Wilcoxon rank-sum test following frequency count normalization. MKMC’s DE results showed substantial overlap and statistically significant concordance with gene-level DEGs identified by the STAR + featureCounts pipeline. In addition, MKMC identified genes not detected by this gene-level workflow, suggesting that k-mer–based representations can capture signals that may be attenuated by gene-level aggregation. However, this comparison reflects differences relative to a gene-count–based pipeline rather than a full transcript-or isoform-aware framework, and therefore should be interpreted within that context. The aggregation of significant k-mers to gene-level signals represents an important methodological consideration. In the current implementation, a gene is labeled as significant if at least one associated k-mer is significant. This approach may bias detection toward longer genes or genes with higher k-mer coverage, complicating direct comparisons with gene-level differential expression results. Future work will explore more principled aggregation strategies that account for gene length, k-mer density, and statistical dependence among features. To assess biological relevance, we performed HCR-based validation of a candidate MKMC-specific signal corresponding to an alternative epha3 isoform. While the observed expression patterns are consistent with differential regulation, the limited number of biological replicates and the nested structure of cell-level measurements within animals constrain statistical inference. As such, these results provide preliminary support for the biological relevance of MKMC-derived signals but require further validation in larger cohorts. We also evaluated MKMC’s performance in transcriptomic age prediction. Using LOOCV and Elastic Net regression, models based on k-mer features achieved performance comparable to gene-based models in this dataset. However, given the small sample size and high dimensionality of the feature space, these results should be interpreted as a proof of concept demonstrating the feasibility of k-mer–based predictive modeling rather than as a definitive assessment of predictive accuracy. While MKMC performs competitively with alignment-based approaches, its reliance on mapping k-mers back to reference annotations for interpretation introduces ambiguity when multiple transcripts share sequence similarity. Ongoing development of de novo assembly and transcript reconstruction modules within MKMC will help address this limitation and improve interpretability. Finally, although MKMC is motivated in part by applications to non-model organisms, the analyses presented here were performed in a system with an available reference genome. Nonetheless, the ability of MKMC to operate without alignment suggests that it may be particularly valuable in settings where reference genomes or annotations are incomplete. In summary, MKMC provides a scalable and flexible framework for k-mer–based transcriptomic analysis. By capturing sequence-level variation directly, MKMC complements existing RNA-seq workflows and can recover biologically meaningful signals that may not be readily detected using standard gene-level pipelines.

**Table 2:**
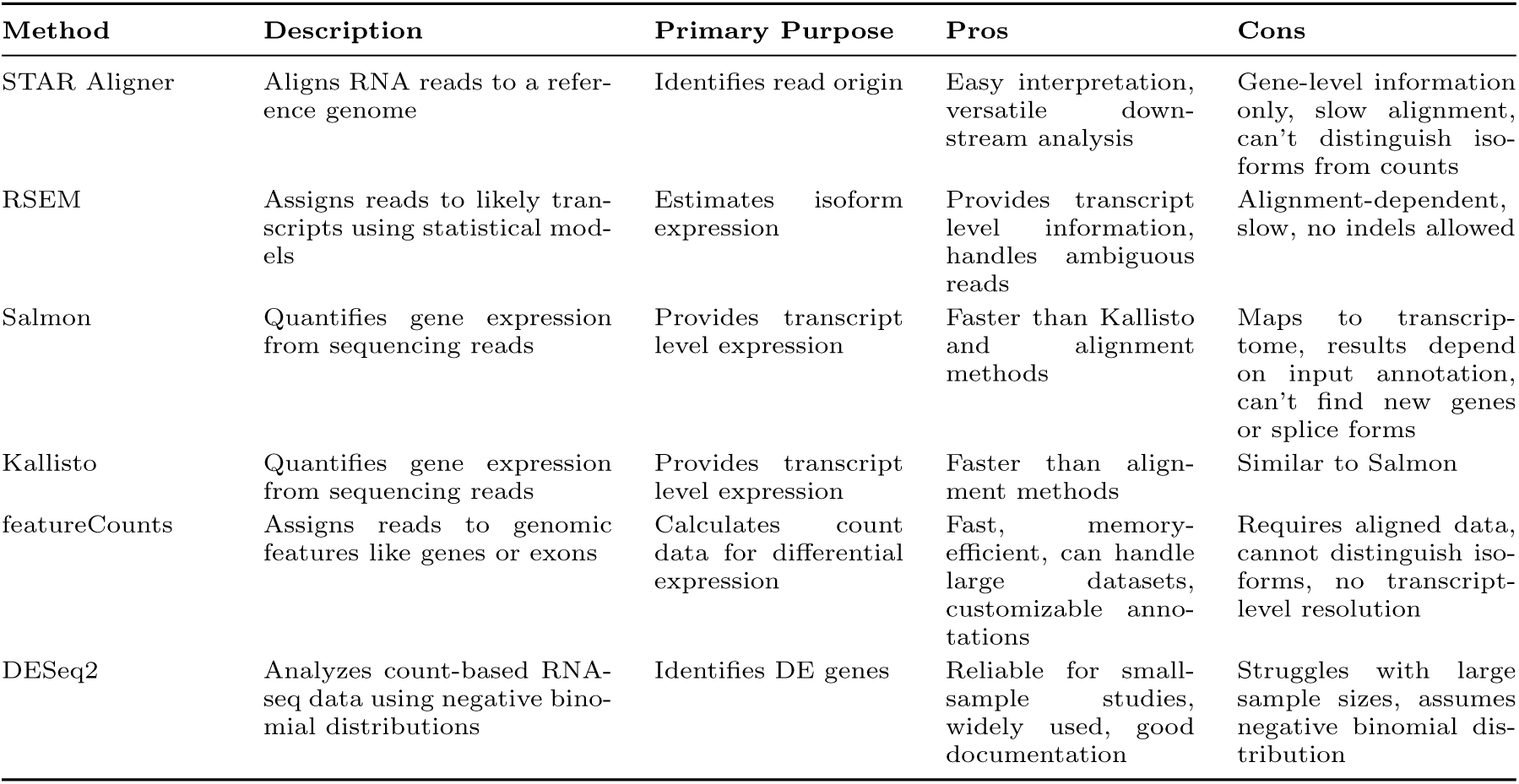
Comparison of common bioinformatics tools for RNA-seq analysis.

**Table 3:** Representative k-mer based methods for reference-free biological discovery and analysis.

| Method: Year | Description | K-mer Counter | Limitations |
| --- | --- | --- | --- |
| **Jellyfish: 2011** | Efficient tool for counting k-mers in large sequencing datasets. | Jellyfish | Slow for larger datasets and memory intensive. |
| **Kissplice: 2012** | Algorithm to extract alternative splicing events, SNPs, and approximate tandem repeats using De Bruijn graphs. | No separate k-mer counter method | Slow on larger datasets. |
| **KOVER: 2016** | Identifies genomic biomarkers using a k-mer representation of genomes and machine learning. | Ray Surveyor Tool | Segmentation faults on some datasets, limited classifier performance. |
| **DE-kupl: 2017** | Captures local RNA variation in RNA-seq libraries, independent of a reference genome. | Jellyfish | Tied to Jellyfish for k-mer counting. |
| **HAWK: 2018** | Association mapping using k-mers. | Modified Jellyfish | Segmentation faults, limited to specific cases. |
| **GECKO: 2019** | Genetic algorithm to classify and explore sequencing data. | Jellyfish | Slow on large datasets. |
| **iMOKA: 2020** | Feature reduction using Naive Bayes classifier with an adaptive entropy filter. | KMC3 | Computationally cumbersome and slow. |
| **Kmerator: 2021** | Extracts specific k-mer signatures and quantifies them into RNA-seq datasets. | countTags | Requires predefined k-mers; geared toward the human genome. |
| **Kmtricks: 2022** | Constructs Bloom filters for large sequencing data collections. | kmtricks | Only generates k-mer abundance matrices. |
| **Kmdiff: 2022** | Differential k-mer analysis for GWAS. | kmtricks | Geared towards GWAS and two-condition studies. Assumes data follows a poisson distribution. |
| **SPLASH: 2023** | Statistical framework for reference-free discovery of biologically relevant k-mers directly from sequencing data across multiple experimental designs and omics applications. | contained netflow pipeline that includes k-mer counting | Primarily focused on identifying significant sequence signatures rather than generating integrated k-mer abundance matrices for downstream transcriptomic analyses. |
| **KaMRaT: 2024** | Processes large k-mer count tables for RNA-seq data with various scoring and filtering tools. | DE-kupl JoinCounts & Jellyfish | Limited to $k \leq 32$. |
| **CellSpace: 2024** | Embeds DNA k-mers and cells into a shared space to integrate scATAC-seq datasets. | Internally optimized method | Designed for scATAC-seq data, not applicable to RNA-seq directly. Limited to $k \leq 10$ which does not map back well to reference genomes. |
| **SPLASH2: 2024** | Detects regulated sequence variation in massive datasets, including splicing and circular RNA. | KMC | Focused on variation detection; less suited for transcript-level quantification. |
| **sc-SPLASH: 2025** | Extends SPLASH for single-cell sequencing to detect splicing, mutations, and novel features. | BKC (Barcoded k-mer Counting) | Primarily for scRNA-seq and spatial data; not generalizable to all omics. |

**Table 4:**
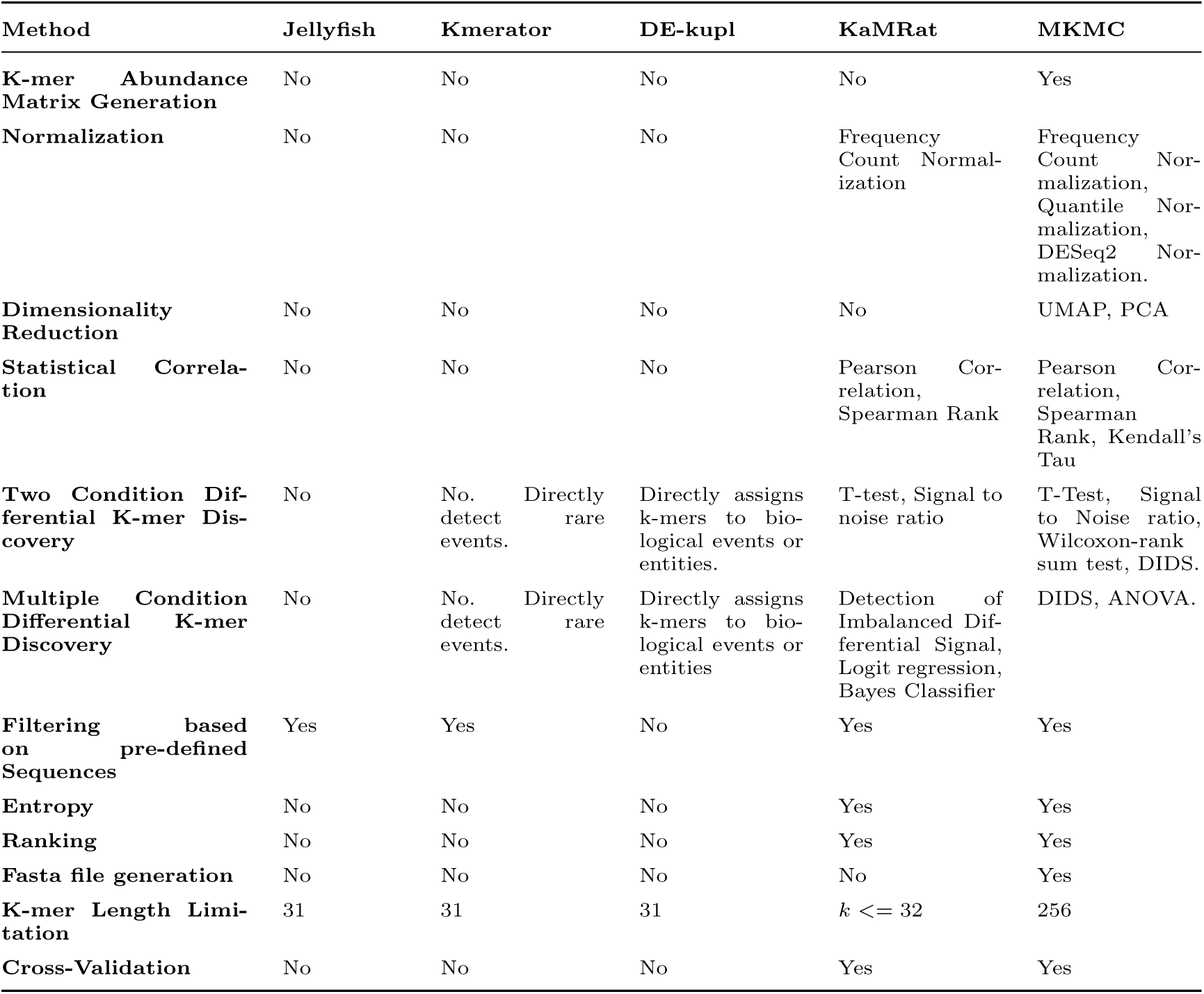
Some of the state of the art k-mer based methods specific to RNA-seq data.

## 5 Materials and Methods

### 5.1 African turquoise killifish husbandry

We used the GRZ strain of the African turquoise killifish species *Nothobranchius furzeri* for the validation experiment. The fish were housed in the Stanford Research Animal Facility under protocols approved by the Stanford Administrative Panel on Laboratory Animal Care (IACUC protocol #13645) under the following conditions: the circulating water system was set to 26 °C, with a conductivity between 3500 and 4500 µS/cm, a pH between 6.8 and 7.2, and a daily exchange of 10% with water treated by reverse osmosis. A 12-hour day/night cycle was used.

Our validation fish were raised as follows. First, fish embryos were collected from group breeding tanks (one male and three females in a 9.8 L tank) or individual breeding tanks (one male and one female in a 2.8 L tank), where the parents were bred with sand trays for 4 hours. Embryos were separated from the sand by gently sieving [42], rinsed twice in 4 mL embryo solution (Ringer’s solution [Sigma-Aldrich, 96724] supplemented with 0.01% methylene blue [Kordo, #37344]), and placed in fresh embryo solution in 30 mm × 15 mm Petri dishes (E Greiner Bio-One, 627102) at a density between 10 and 30 embryos per plate and incubated at 28 °C for 2 weeks. Next, the embryos were placed on moist coconut fiber at 27 °C to incubate for 2 weeks, and then fish were hatched in 10 mL chilled (4 °C) 1 g/L humic acid solution (Sigma-Aldrich, 53680) and incubated at room temperature overnight. The hatched fish were transferred to the animal facility and placed in 0.8 L tanks at a density of 10–20 fish per tank, split to a density of 4 fish per 0.8 L tank in the second week after hatching, 2 fish per 0.8 L tank in the third week, and one fish per 0.8 L tank in the fourth week. Live brine shrimp was used to feed the juvenile fish daily until sexual maturity. Five weeks after hatching, each fish was sexed by the tail fin color (males and not females exhibit vivid colors), randomized, and transferred to a 2.8 L tank. Fish with severe gill defects, curved spines, and an inability to float (“belly sliders”) were excluded. Once the fish were housed in 2.8 L tanks, they were fed using a custom-made feeder twice a day, 15 mg per feeding, for 30 mg of Otohime fish pellet (Reed Mariculture, Otohime C1) daily. To provide enrichment for the fish, we placed a sand tray in each tank for 4 hours per day, two days per week. This enrichment leads to some spontaneous release of eggs in females, and the females do not exhibit unhealthy “egg-bound” phenotypes characterized by abdominal swelling during lifetime.

### 5.2 Sample collection

All procedures were performed in an RNAse-free environment, with nuclease-free tools and reagents. Fish were randomly assigned a harvest order and harvested at 2 months of age. On the day of harvest, each fish was fed 15 mg of Otohime fish pellet 8:30-8:45 AM and typically sacrificed from 11:30 AM to 1:30 PM (3-5 hours after the last meal). The fish were sacrificed in ice slurry (0 °C) for 2-3 min, dried, and placed on an ice-cold Sylgard-coated Petri dish filled with wet ice and covered in plastic wrap. The skin covering the gills and the abdominal muscles on both sides of the fish were removed to expose the internal organs. The dissected fish were incubated in 6-10 mL 4% PFA in PBS (Santa Cruz, CAS 20525-89-4) on a rocking platform at 4 °C for 20-24 hours, washed 3 times in 12 mL PBS on ice (1 hour incubation for each wash), placed in 12 mL fresh PBS to store at 4 °C until dissection (up to 4 days). The fish were dissected in ice-cold PBS, and the livers were incubated on ice (10 min each) in 1 mL 66% methanol:33% PBS, 1 mL methanol, 1 mL methanol, and stored in 1 mL methanol at-20 °C until sample sectioning. A blinding ID was assigned to each sample, and this ID was used for the subsequence procedures until after image quantification.

To prepare for cryosectioning, the livers were rehydrated by incubating (10 min each) in 1 mL 75% methanol:25% PBS, 1 mL 50% methanol:50% PBS, 1 mL 25% methanol:75% PBS, and 2 times in 1 mL PBS at room temperature. Next, the tissues were incubated in 1 mL nuclease-free 30% sucrose solution (30 g sucrose in 100 mL PBS) at 4 °C for 20-24 hours. For embedding, each liver was dried using Kimwipe (Fisher Scientific, 06-666A), placed in a Peel-A-Way histology mold (Ted Pella, 27110) filled with optimal cutting temperature (OCT) compound (Fisher Scientific 23-730-571) along the anterior-posterior axis with the concave side down. The OCT blocks were frozen in ethanol-dry ice bath for 15 min and then stored at-80 °C. Each sample was sectioned at a 10-25 µm thickness using a MX35 Ultra microtome blade (Epredia, 3053835) in a cryostat (Leica, CM3050 S) at-18-20 °C objective and chamber temperatures. All the tissue sections were captured on the Superflost Plus slides (Fisher Scientific, 12-550-15), dried at temperature for 15 min, and stored at-20 °C. The experimental metadata are listed in Supplemental File 1.

### 5.3 Hybridization chain reaction (HCR) for RNA in situ hybridization

#### 5.3.1 HCR probe design

All the HCR probe sequences are listed in Supplemental File 2. We designed HCR probes using a custom-made Python script *Probe Maker.py* [43]. To detect an mRNA molecule, HCR uses an ensemble of 20–50 sequences that anneal to the mRNA, and this ensemble is referred to as a ‘probe set.’ Each HCR sequence has a bi-partial feature, with half of the sequence corresponding to a specific HCR initiator sequence (e.g., B1h1, B1h2, etc.) and the other half corresponding to the transcript-specific sequences (see Choi et al., 2018 for details). The common probe set of the *epha3* gene (*epha3-com*) includes 42 sequences and corresponds to exons 8–15 of the *epha3* gene, spanning a ∼1000-bp region not detected as ‘male specific’ by MKMC. The male-specific probe set of *epha3* (*epha3-spe*) includes 24 sequences, corresponding to the 571-bp sequence (contained in exon 4) detected as ‘male specific’ by MKMC. Both probe sets were purchased from IDT as an oPool Oligo pools (50 pmol, then resuspended to 0.5 pmol/*µ*l in RNase-free TE 8.0 [Ambion, AM9858]). All the HCR reagents were purchased from Molecular Instruments (HCR amplifiers), with the B5 hairpins conjugated to Alexa Fluor 647 and the B3 hairpins to Alexa Fluor 546.

#### 5.3.2 Staining Procedure

All the procedures were conducted in an RNase-free condition. The liver sections were washed twice with 1 mL PBS for 5-minute washes, and then four times with 500 *µ*L PBST for 5-minute washes at room temperature. Next, the slides were pre-hybridized using 150 *µ*L of the hybridization buffer (Molecular Instruments, HCR tissue probe hybridization buffer) at 37 ^◦^C in a humidified chamber for 30 min to 1 hour. The common and male-specific probe sets were mixed with the hybridization buffer (add 1 *µ*L of each 0.5 pmol/*µ*L probe in 100 *µ*L hybridization buffer), and ∼80 *µ*L of the mixture was added to each slide (a coverslip was added on each slide to reduce evaporation) and incubated at 37 ^◦^C in a humidified chamber for 19–20 hours. The slides were washed with probe wash buffer (Molecular Instruments, HCR tissue probe wash buffer), 15 min each, in the following solutions: 500 *µ*L 100% Wash Buffer, 500 *µ*L 75% Wash Buffer:25% 5× SSCT (0.1% Tween, diluted from 20× SSC [Invitrogen, AM9770] in nuclease-free water), 500 *µ*L 50% Wash Buffer:50% 5× SSCT, 500 *µ*L 25% Wash Buffer:75% 5× SSCT, and 500 *µ*L 5× SSCT. The slides were pre-amplified with ∼150 *µ*L Amplification Buffer (Molecular Instruments, HCR amplification buffer) at room temperature for ∼1 hour. Hairpin solutions were prepared per the manufacturer’s instructions (each hairpin was incubated at 95 ^◦^C for 90 sec, then cooled down at room temperature in the dark for ∼25 min). Amplification solution was prepared by mixing 2 *µ*L of each hairpin (3 *µ*M) into 100 *µ*L Amplification Buffer. About 50 *µ*L of the amplification solution was incubated with the slides at room temperature for ∼20 hours. The amplification solution was removed and the slides were washed twice (30-minute incubation per wash) with 500 *µ*L DAPI/5× SSCT solution (0.01 mg/mL DAPI diluted in 5× SSCT). The slides were mounted using antifade reagent (Thermo Fisher Scientific, ProLong Gold Antifade Mountant, P36934) and 22 × 50 mm, No. 1.5 coverslips (Fisher Scientific, 12-544-DP), and sealed with nail polish.

#### 5.3.3 Confocal microscopy

We imaged each slide as 16-bit images on an inverted confocal microscope (LSM 900), equipped with ZEN Blue software (3.0) and a 40× objective (ZEISS Plan/Apochromat/40×/1.40, oil) and Zeiss Immersol oil 518F (Zeiss, 4Y00-R0DY-1007-3VF3). The images were acquired as *z*-stacks, with 15 *z*-slices and a 0.5 *µ*m step size. Using a pinhole size of 32 *µ*m, we imaged three channels, including Alexa 546 (577/603; 1.5% laser intensity, detector gain: 775 V, detector offset: 256, detector digital gain: 1.0); DAPI (353/465; 0.5% laser intensity, detector gain: 650 V, detector offset: 256, detector digital gain: 1.0); and Alexa 647 (653/668; 8.0% laser intensity, detector gain: 650 V, detector offset: 512, detector digital gain: 1.0). Each *z*-slice was imaged in all three channels before moving to the next *z*-slice. We imaged two evenly spaced fields of view on each liver section.

#### 5.3.4 Image analysis and quantification

Each image was first maximum-intensity projected in the z-direction using a custom-made FIJI macro script (FIJI, 2.9.0) [44]. Max-projected images were loaded into QuPath software (v.0.5.1, https://qupath.github.io/). The cells were segmented using a nuclear mask created based on the DAPI signal (DAPI threshold: 2000, sigma: 1.5, minAreaMicrons: 10.0, maxAreaMicrons: 400.0, backgroundRadiusMicrons: 8.0, and other default parameters) and then an expansion 3 µm from the DAPI mask was used as the cell boundaries. Detection of red blood cells, which have strong autofluorescence in all channels, was manually removed to avoid false positive subcellular spot detection. Next, the QuPath subcellular detection function was used to detect each type of mRNAs using specific parameters (see Supplemental File 3 for details). Every cell was visually inspected to check the detected spots matched with manual counting. A small number of false positive spots were manually removed (these spots usually occur in regions overlapping with red blood cells). The QuPath quantification was exported as.csv files using the ‘Cells’ or ‘Detections’ setting (Supplemental Files 3 and 4).

Summary statistics were calculated as follows. The number of a specific mRNA type per cell and the total cell number of each image was used for subsequent calculations. For the figure related to the expression distributions (e.g., Figure 4C), the cells from all four animals were pooled for a given biological sex (male or female). The Kolmogorov-Smirnov test was used to compare between the distributions. To calculate the summary statistics of each animal, the cells from the two fields of view were pooled. The average number of a specific mRNA type per cell was calculated by dividing the total number of this given mRNA by the total number of cells using Excel. The ratio of the common probe spots (*epha3-com*) the male-specific probe spots (*epha3-spe*) was calculated by dividing the average number of ‘epha3-com’ spots by that of ‘*epha3-spe*’ spots. After completing all the calculations, the sample information was ‘de-blinded,’ and the biological sex of each sample was revealed. The Mann-Whitney test was used to assess statistical significance. Data plotting and statistics were performed in Prism 10 (GraphPad).

Colocalization analysis was performed as follows. To determine whether two spots colocalized, we first identified a distance threshold (in µm) that naturally separates the distances between spots that are very close to each other from those farther apart. We calculated the distances between all the spot pairs in every cell. For example, for a cell with ‘n’ spots in the Ch1 channel (‘*epha3-spe*’) and ‘k’ spots in the Ch4 channel (‘*epha3-com*’), the number of spot pairs in this cell would be the product of *n* and *k*. We filtered out the cells with no spots detected in either fluorescence channel. The distance between each spot pair was calculated as the square root of (*x*_1_ − *x*_2_)^2^ +(*y*_1_ − *y*_2_)^2^, where (*x*_1_*, y*_1_) is the centroid coordinate of a Ch1 channel spot and (*x*_2_*, y*_2_) is the centroid coordinate of a Ch4 channel spot. The centroid coordinates of every spot were exported from the QuPath quantification. We plotted the distance distribution of all spot pairs, which forms a bimodal distribution with a separation at 0.375 µm (Figure 4F). If the centroids of two spots are less than 0.375 µm from each other, then these two spots would be called ‘colocalized.’ In contrast, two spots would be considered as ‘non-colocalized’ if they were farther than 0.375 µm from one another. Lastly, we calculated the number of colocalized spots in each cell as the number of spot pairs with distances *<* 0.375 µm. The number of ‘*epha3-spe*’-only spots in a cell was calculated by subtracting the number of colocalized spots from the total number of ‘*epha3-spe*’ spots in that cell, and the number of ‘*epha3-com*’-only spots was calculated similarly, except using the total number of ‘*epha3-com*’ spots in that cell. In 5 cells (out of *>* 2000 cells analyzed), a single ‘*epha3-spe*’ spot was found to be near more than one ‘*epha3-com*’ spot, leading to a ‘-1’ count for ‘*epha3-spe*’-only or ‘*epha3-com*’-only spots. These cells were excluded from the analysis. Summary statistics and the statistical significance comparing different conditions for the colocalized spots, ‘*epha3-spe*’-only spots, and ‘*epha3-com*’-only spots were calculated as above. The full data can be found in Supplemental Files 5 and 6.

### 5.4 Bulk RNA-sequencing of Killifish Retina-RPE

#### 5.4.1 Retina-RPE Dissection

Animals were raised as described in Section 4.1, with one exception: all of the animals used for this study were the product of multiple generations of backcrossing a mutant line with the Nothobranchius furzeri GRZ strain. Adult male fish were euthanized using a slurry of ice and system water and placed in a 60 x 15 mm dish (Greiner Bio-One, 628161) containing ice-cold 1X phosphate-buffered saline (PBS; diluted from 10X PBS [Alfa Aesar, J62036-K2] using nuclease free water). Using spring scissors (Fine Science Tools, 91500-09), the head was bisected. To extract the eyes, micro scissors were gently inserted behind the eye to visualize the optic nerve, after which the eyeball was carefully excised using forceps (Fine Science Tools, 11251-20) and scissors. The nasal caruncle served as a landmark during dissection. Isolated eyeballs were transferred to fresh cold 1X PBS in a 60 x 15 mm dish using a transfer pipette (Fisher Scientific, 13-711-20). While in PBS, extraocular muscles were removed, and the optic nerve was severed with scissors. A small incision was made in the cornea using spring scissors, allowing the cornea to be gripped with forceps and peeled back to expose the lens, which was then removed. The iris was then cut circumferentially with scissors, and the sclera was carefully removed. The choroidal layer was separated away using fine forceps. The RPE was kept intact to minimize damage to the underlying retina. The resulting retina-RPE sample was washed thoroughly in 1X PBS, snap-frozen in a 1.5 mL tube (CELLTREAT, 229443) in liquid nitrogen, and stored at-80°C until RNA extraction. This process was repeated for each sample.

#### 5.4.2 RNA Isolation

RNA extraction was performed using the QIAGEN RNeasy Plus Micro kit (QIAGEN, 74034), according to the manufacturer’s instructions. RNA was eluted in a final volume of 13 µL.

#### 5.4.3 Bulk RNA-Sequencing Library Preparation & Sequencing

The sequencing library was prepared and sequenced by GENEWIZ/Azenta (Azenta, South Plainfield, New Jersey, United States) using the Nextera XT DNA Library Preparation Kit (24 samples; Illumina, FC-131-1024) and sequenced on one lane of a NovaSeqS4. GENEWIZ/Azenta performed base calling, demultiplexing, and FASTQ file generation.

| Label | Group | RIN | RNA con. (ng/μl) | Total RNA con. (ng/13μl) |
| --- | --- | --- | --- | --- |
| BC1 | Male_1.5 mo | 10.0 | 47 | 611 |
| BC2 | Male_1.5 mo | 10.0 | 36 | 468 |
| BC3 | Male_1.5 mo | 10.0 | 23 | 299 |
| BC4 | Male_3.0 mo | 9.5 | 15 | 195 |
| BC5 | Male_3.0 mo | 9.6 | 10 | 130 |
| BC6 | Male_3.0 mo | 9.9 | 72 | 936 |
| BC8 | Male_4.5 mo | 10.0 | 5 | 65 |
| Resub-BC13 | Male_4.5 mo | 9.8 | 80 | 1040 |
| Resub-BC14 | Male_4.5 mo | 7.9 | 104 | 1352 |
| BC10 | Male_6.0 mo | 9.7 | 21 | 273 |
| BC11 | Male_6.0 mo | 9.4 | 8 | 104 |
| BC12 | Male_6.0 mo | 9.4 | 86 | 1118 |

### 5.5 Read Mapping using STAR

Raw sequencing data (FASTQ files) for the liver were downloaded from Run Selector and checked for quality and trimmed using Trim-galore v0.5.0. The processed paired-end RNA-seq reads were aligned to the African turquoise killifish reference genome downloaded from NCBI (Nfu 20140520, GCF 001465895.1) using STAR v2.7.10b with the default parameters. The genome index, pre-generated with splice junction annotations to optimize spliced read alignment, was used. During alignment, STAR handled on-the-fly decompression of.fq.gz files, and output BAM files were sorted by genomic coordinates. Default parameters were employed to allow mismatches (up to 10 or 30% of the read length) and to permit insertions and deletions (indels), though with penalties applied to minimize their occurrence. Gapped alignments spanning introns were supported with default intron size thresholds (21–1,000,000 bases). Reads mapping to more than 10 loci were excluded from the final alignments, ensuring high-confidence mappings. Futhermore, Samtools v1.16.1, with the parameters of MAPQ *<* 255 (‘samtools view-q255-b’), was used to remove the reads mapped to multiple genomic regions. Next, we inputted the uniquely mapped reads into the ‘featureCounts’ program (with the default parameters) from subread v2.0.6 to generate the read counts for each gene.

### 5.6 Dimensionality Reduction K-mer Visualization

To reduce dimensionality reduction of k-mers matrix we used MKMC with the command below:

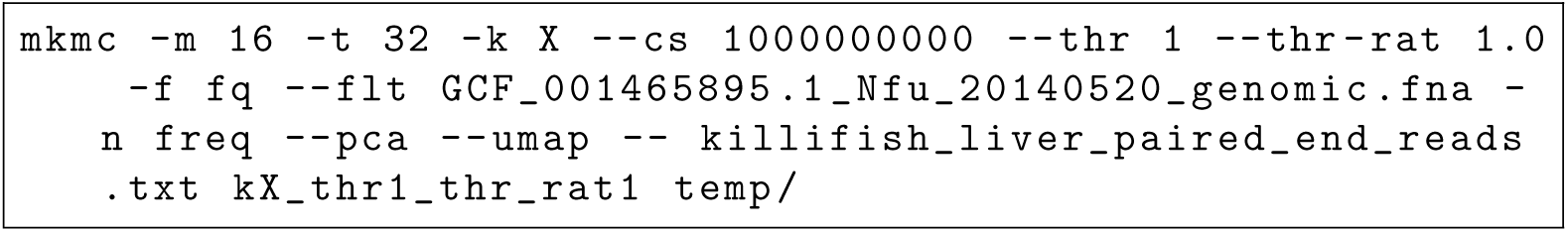

We generated frequency-normalized matrices for canonical k-mers of length X (10 to 50 in steps of 20) with at least one count in all the samples by filtering only the k-mer from the killifish genome. The matrices were the base to perform MKMC built-in Principal Component Analysis (PCA) and Uniform Manifold Approximation and Projection (UMAP). PCA reduces dimensionality by identifying orthogonal components that capture the maximum variance in the data, while UMAP is a non-linear technique that preserves both the global and local structure of high-dimensional data.

### 5.7 Differential Expression Analysis of K-mers using MKMC

The command line below was used to conduct the k-mer (k=30, 50) differential expression analysis:

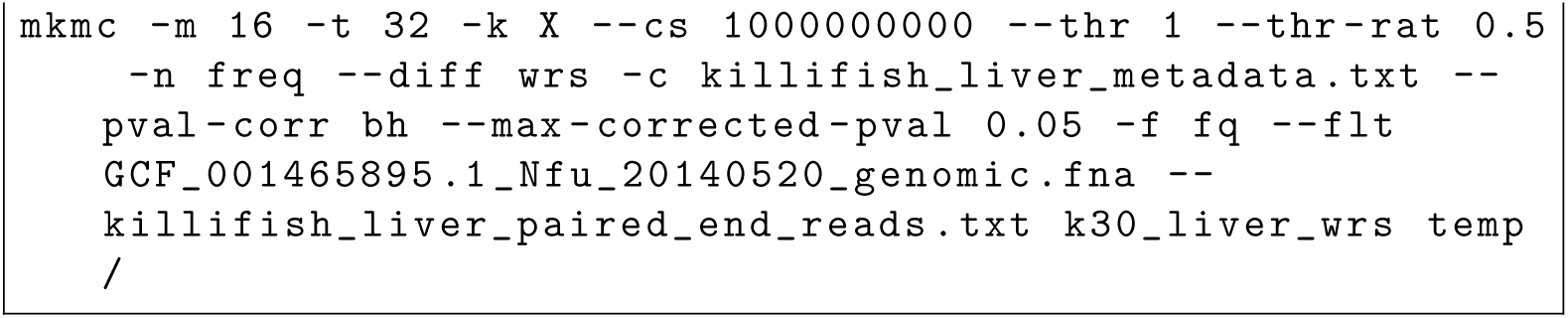

This command line generated canonical 30-mer or 50-mer binary matrices from the paired-end reads with at least 1 count in half the samples. Furthermore, it uses frequency count normalization to normalize the k-mers per samples before running the wilcoxon rank sum test adjusted by benjamini-hochberg method. Finally, it filters out k-mers to only output those with a p-value below 0.05 (FDR threshold). All the k-mers used are guaranteed to only be coming from the reference genome when using the --flt parameter.

Once the differentially expressed k-mers are identified, they are outputted as a separated fasta file by MKMC which can be used to align them to the reference genome using any alignment tool to identify the genes they originated from. For example, we used BWA-MEM to align the k-mers with the bash command below:

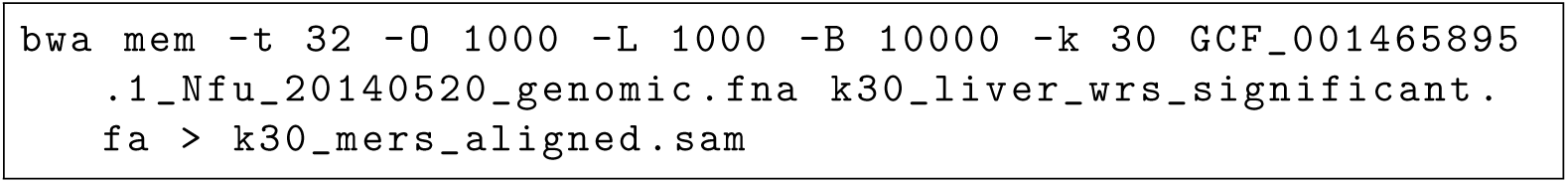

Here, we do not allow indels and matches the k-mers with 100% accuracy. Next, we used the bam file generated by BWA-MEM k-mer alignment to generate the k-mer counts for each gene using the ‘featureCounts’ program (with the default parameters) from subread v2.0.6:

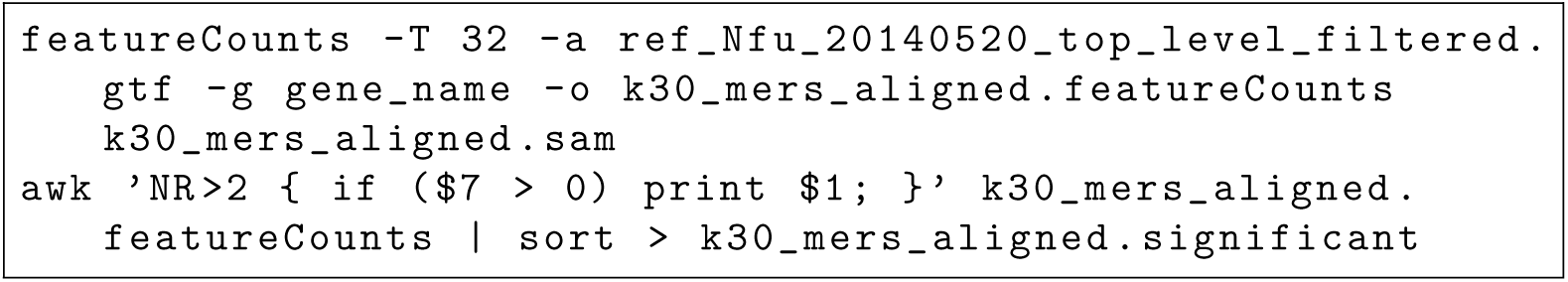

### 5.8 Comparison of DE K-mers Genes and Traditional Alignment

To compare the overlap between differentially expressed (DE) genes identified by the two methods—k-mer-based analysis and alignment-based analysis—ranked gene lists were generated for each method. Each list was converted into rank positions, and the overlap of top-ranked elements between the lists was analyzed at varying thresholds (e.g., top 100, 200, etc.).

For each pair of thresholds, the overlap between the top-ranked genes was computed and tested for statistical significance using the hypergeometric test. The total number of genes in the ranked lists (*M*), the number of genes in the top subsets (*n* and *N*), and the observed overlap (*k*) were used to calculate p-values via the survival function of the hypergeometric distribution. The resulting p-values were adjusted for multiple hypothesis testing using the benjamini-hochberg procedure. Finally, the adjusted p-values were thresholded so that any p-value below 0.05 (FDR threshold) would be significant (1) and any value above would mean the overlap is not significant.

The heatmap was generated to display the significance across the combination of overlaps between top gene sets from the two methods. This approach highlights the concordance or divergence in identifying DE genes between the k-mer and alignment-based methods.

### 5.9 K-mer Age Prediction using Elastic Net Regression

Age prediction was performed using scikit-learn in Python. The *k*-mer count matrix was generated using MKMC and subsequently log transformed to stabilize the variance and then standardized using StandardScaler from sklearn.preprocessing. To construct the matrix, only canonical *30*-mers with at least 500 counts across all samples were retained using the following command:

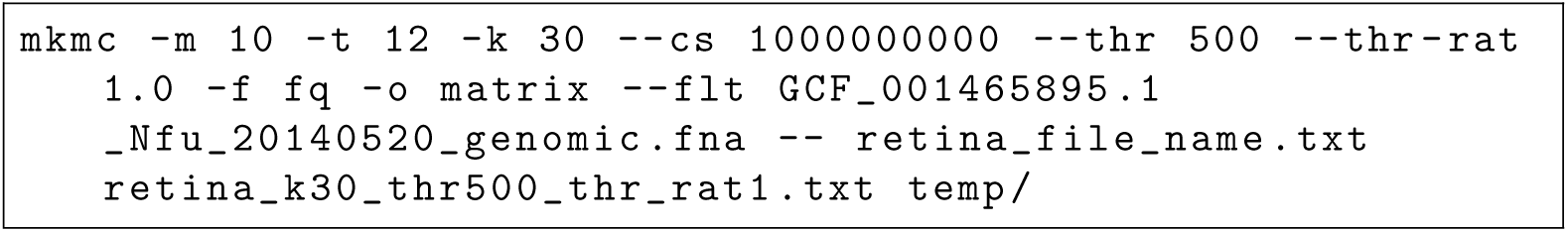

Elastic Net regression (sklearn.linear model.ElasticNet) was used for predictive modeling, with Leave-One-Out Cross-Validation (LOOCV) (sklearn.model selection.LeaveOneOut) employed to evaluate performance. The regularization parameters (*α* and *l1 ratio*) were optimized through grid search over predefined values (*α* = [0.001, 0.01, 0.1, 1, 10] and *l1 ratio* = [0.0, 0.1,…,1.0]), selecting the combination that minimized the Mean Absolute Error (MAE) (sklearn.metrics.mean absolute error).

Model performance was evaluated using MAE, R-squared (*R*^2^) (sklearn.metrics.r2 score), and Pearson correlation (*r*) (scipy.stats.pearsonr).

## 7 Code Availability

The MKMC code can be downloaded from GitHub: https://github.com/refresh-bio/MKMC.

The Bash, Python, R scripts and any additional images and files used for the analysis of this paper, if not presented in supplementary material, can be found at: https://github.com/lajoycemboning/MKMC analysis

## 8 Data Availability

The african turquoise killifish liver tissue used in this study is deposited in the Gene Expression Omnibus (GEO) database with the accession number: GSE216369. The additional Tabula Muris Senis dataset used to benchmark MKMC performance is deposited with the accession number: GSE132040. The retina samples used in this study is deposited in the GEO database with the accession number: GSEXXXXXX.

## 9 Author Contributions

## Supporting information

Retina Counts

Retina Normalized Counts

## 10 Acknowledgements

We thank Megan Mitchell for her early testing of MKMC and for providing valuable feedback that helped improve the toolkit. We also thank Param Priya Singh for insightful discussions on addressing the genomic complexity of the African turquoise killifish.

## 13 Supplementary Material

### 13.1 Details on MKMC functionalities

#### 13.1.1 Normalization

Normalization in MKMC addresses sequencing depth and technical variability to ensure k-mer counts are comparable across samples. The toolkit offers multiple normalization methods to suit diverse datasets and analysis objectives:

- Frequency Count Normalization: Converts raw k-mer counts into relative frequencies for straightforward comparisons.
- Quantile Normalization: Aligns the distributions of k-mer abundances across samples, reducing systematic biases.
- DESeq2-like Normalization: Adapts a size factor approach for k-mer abundance matrices to account for differences in sequencing depth while controlling for outliers.

#### 13.1.2 Dimensionality Reduction

High-dimensional k-mer datasets can be challenging to interpret and computationally expensive to analyze. MKMC provides built-in dimensionality reduction techniques to address these challenges:

- Principal Component Analysis (PCA): Projects k-mer data into orthogonal components that capture the most variation, simplifying exploratory data analysis.
- Uniform Manifold Approximation and Projection (UMAP): Preserves both local and global structure in a low-dimensional space, aiding in visualization and cluster identification.

#### 13.1.3 Statistical Correlations

MKMC includes a comprehensive suite of tools for statistical correlation analysis, enabling users to identify associations between k-mer abundances and experimental conditions. Supported methods include:

- Pearson Correlation: Measures linear relationships between k-mer counts and continuous variables.
- Spearman Rank Correlation: Detects monotonic relationships, including non-linear patterns.
- Kendall’s Tau: A robust method for analyzing ordinal data or datasets with tied ranks.

#### 13.1.4 Two Condition Differential K-mer Discovery

K-mers are selected according to a binary conditions (control vs. disease).

##### T-Test

Firstly a log2(x + 1) transformation is applied to sample counts. Then each feature association between sample counts and labels is evaluated by p-value t-test, adjusted by Bonferroni, Benjamini-Hochberg, Benjamini-Yekutieli or Holm-Bonferroni correction procedure for controlling false discovery rate. The top k-mers are selected from the lowest value to the highest p-values.

##### Signal-to-Noise ratio

The signal to noise ratio (SNR) is a measure used to compare the level of a desired signal to the level of background noise [45]. The signal is the expression levels of k-mers, while the noise is the variability or errors in the measurements. The SNR is calculated by dividing the difference between group means by the sum of group standard deviation. The top k-mers are selected with absolute values from highest to lowest.

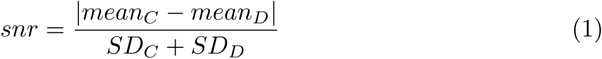

##### Wilcoxon-rank sum test

The normalized k-mer counts are used to conduct the Wilcoxon rank-sum test. Then each feature association between sample counts and groups is evaluated by p-value, adjusted by Bonferroni, Benjamini-Hochberg, Benjamini-Yekutieli or Holm-Bonferroni correction procedure for controlling false discovery rate. The top k-mers are selected from the lowest value to the highest adjusted p-values.

##### Detection of Imbalanced Differential Signal

To detect k-mers with abnormal signal in a multi-condition experiment [46]. For each k-mer, we calculate the maximum expression in the control group.

Let the k-mer expression values of the *n*_1_ control samples for a k-mer be given by:

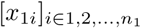

and the k-mer expression values of the *n*_2_ disease samples for a k-mer be given by:

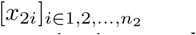

To detect k-mers that show higher expression in control samples compared with disease samples, the maximal k-mer expression value of a k-mer among control samples as:

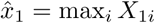

The DIDS score is then given by:

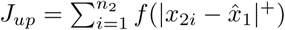

where:

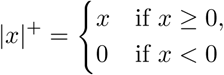

where f is a strictly increasing function. We use the following three variants of f:

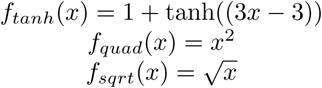

#### 13.1.5 Multiple Condition Differential K-mer Discovery

##### ANOVA

The normalized k-mer counts are subjected to the ANOVA test. This statistical test assesses whether there are significant differences in mean expression levels across multiple groups. Each k-mer feature’s association with group differences is evaluated using ANOVA, generating p-values indicative of significance. P-values are adjusted using Bonferroni, Benjamini-Hochberg, Benjamini-Yekutieli or Holm-Bonferroni correction procedure. Subsequently, k-mers are prioritized based on their adjusted p-values.

##### Detection of Imbalanced Differential Signal

To detect k-mers with abnormal signal in a multi-condition experiment. For each k-mer, we calculate the maximum expression in the control group.

Let the k-mer expression values of the *n*_1_ control samples for a k-mer be given by:

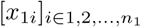

and the k-mer expression values of the *n*_2_ disease samples for a k-mer be given by:

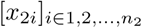

This multiple-condition filtering method returns the maximual value by iteratively estimating the DIDS score by comparing one condition against the other.

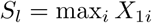

#### 13.1.6 Entropy

We add a +1 to all the counts of the k-mer matrix to avoid flow errors when taking the logarithm [37]. We calculate the entropy of a k-mer across samples using Shannon’s formula. For a given k-mer k and it’s counts in the different samples *S_k_*= (*C_k_*_0_*, C_k_*_1_*,…,C_kn_*), we compute it’s entropy value *H_k_*as follows:

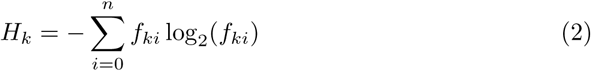

where:

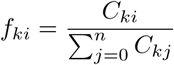

*f* = frequency, *n* is the number of samples. The top *k*-mers are selected from smallest to largest score.

### 13.2 MKMC workflow details

*k*-mers counting process may be parameterised in the aim of deciding which *k*-mers should be processed in the next stages, e.g. for excluding too rare *k*-mers. It is performed simply by passing proper parameters to KMC.

*k*-mers merging (*ii*) is performed independently for each KMC bin. The input is *s* bins, respectively for every sample, containing counts of lexicographically ordered *k*-mers. To merge all k-mers across samples for a given bin a binary heap is employed. The heap is initialized with the first *k*-mers from every sample. The lexographically lowest sequence is on the top of the heap, and it is (if fulfills filtering criteria) the first output. After extracting the top *k*-mer from the heap, the next one of the corresponding sample is added to the heap, unless there are no more *k*-mers in the sample. Then, the same *k*-mer, if any, of other samples is extracted from the heap until the new *k*-mer is found. The single row of a matrix is created with counts of the extracted *k*-mers and zeros for samples that were not present in the heap, meaning that these *k*-mers were absent in such samples. New *k*-mers of the samples are constantly inserted after every extraction. The same process is repeated for the next *k*-mers, until all *k*-mers have been processed.

The rows are saved if they fulfill the given requirements of the fraction of samples with the specified *k*-mer counts. Keeping only *k*-mers belonging to the given additional set is performed by simple *k*-mer counting of sequences in the set. The obtained KMC file is linearly iterated while merging consecutive *k*-mers.

The stage is also responsible for data collection for normalization, e.g. for frequency count, it collects sums of the samples counts to divide the counts by the sum later. The data is stored in temporary binary STATS files.

Depending on the work scenario, MKMC performs a few passes of the input data or its modified form. Generally, every of the abovementioned stages perform a single pass, except correction (*iv*), which performs two passes: correction itself and storing the results.

### 13.3 Details on MKMC performance

We observed the time of various main MKMC stages depending on k-mer length and a number of threads (supplementary Fig. 9). The results include sequential dimensionality reduction resulting in long postprocessing.

We observed CPU utilization (32 threads) and memory usage for performing all the tasks in a single run, except dimensionality reduction (supplementary Fig. 10; similarly to Fig. 3g).

### 13.4 MKMC performance benchmarking experiments running

In the experiments whose results was shown on Fig. 3 and supplementary Fig. 9, 10, we used the datasets given in Table 5 and killfish (for GSE216369) genome stated in the article. We excluded the following samples from GSE132040 due to a lack of a phenotype required in some experiments. SRR9126736, SRR9126839, SRR9126882, SRR9127209, SRR9127374, SRR9127440, SRR9127538, SRR9126793, SRR9126847, SRR9126981, SRR9127233, SRR9127413, SRR9127473, SRR9127564.

**Fig. 10:**
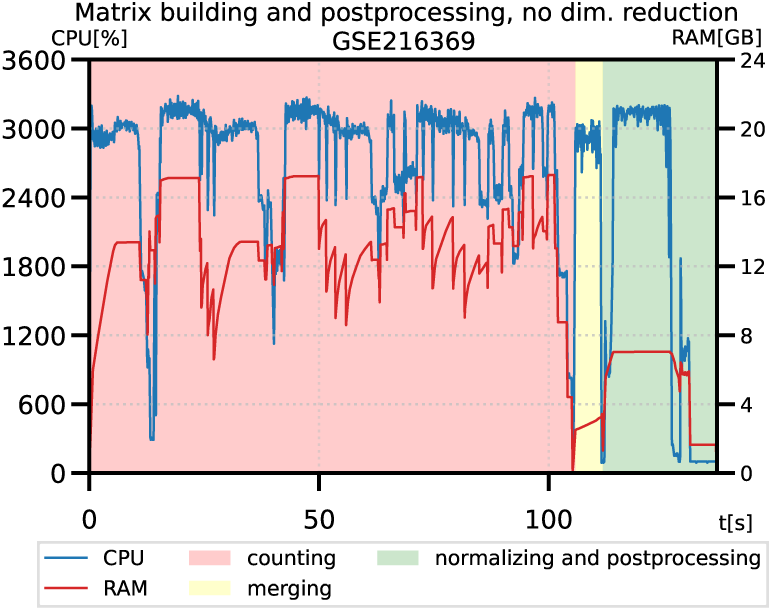
**CPU utilization (32 threads) and memory usage by MKMC performing all the tasks, except dimensionality reduction.**

**Table 5:** Datasets utilized in experiments.

| Dataset | Number of samples | Raw data size | Std. deviation of a single sample size | Number of reads |
| --- | --- | --- | --- | --- |
| GSE216369 | 16 | 224.28 GB | 1.613 GB | 2.45G |
| GSE132040 | 933 | 12.49 TB | 4.689 GB | 177.2G |

Versions of the utilized tools were as follows:

- k-mer processing tools: MKMC — 1.0.0; kmtricks — 1.4.0; Kamrat — 1.2.0;
- mappers: BWA — 0.7.18; STAR — 2.7.11b;
- supporting tools: Jellyfish — 2.3.1; joinCounts has no version; featureCounts — 2.0.8; TrimGalore — 0.6.10;
- system-wide tools: apptainer — 1.4.5; Python — 3.13.9.

To measure wall-clock times and peak memory consumptions we utilized GNU time tool. CPU usage and memory consumption profiles (Fig. 3g and supplementary Fig. 9) were generated with Python cprof tool available in the GitHub repository. All the experiments were repeated once.

#### 13.4.1 Command lines

Main time and memory benchmarks (subplots of Fig. 3 and, when stated, supplementary Fig. 9, 10) show results obtained by running MKMC, kmtricks, and Kamrat with the following command lines. MKMC and kmtricks generated binary matrices only. Kamrat indexed matrix in the text form, which was generated by merging with joinCounts results of Jellyfish dump.

##### MKMC — building a matrix (a, b, c, f, h)

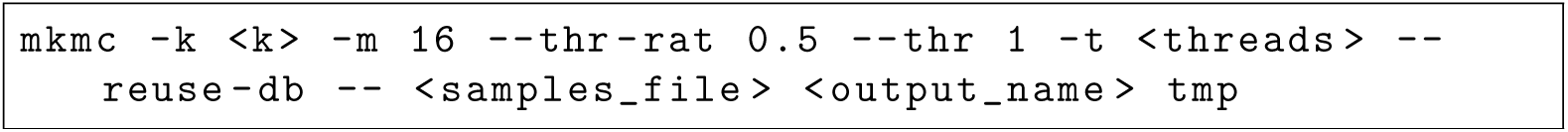

The binary matrix (.kmcdb and.stats files) has not existed before the running. The running also collects data for frequency normalization.

##### MKMC — postprocessing counts by determining correlation (d, e)

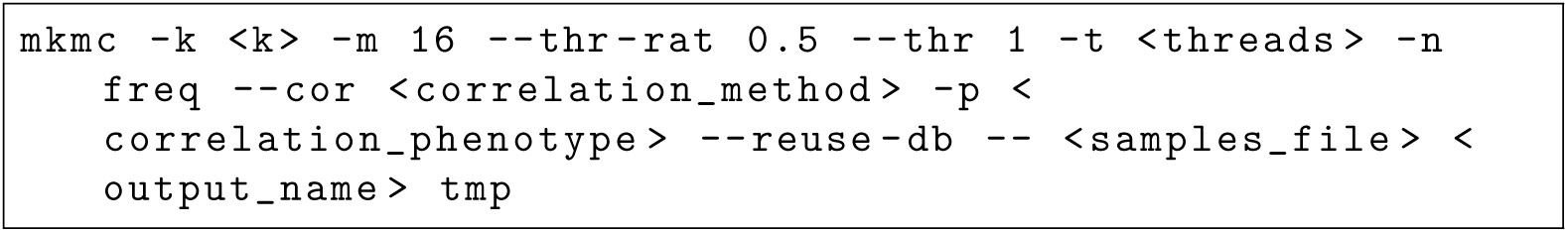

The binary matrix has been generated before the running.

##### MKMC — postprocessing counts by differential k-mers analysis (d, e)

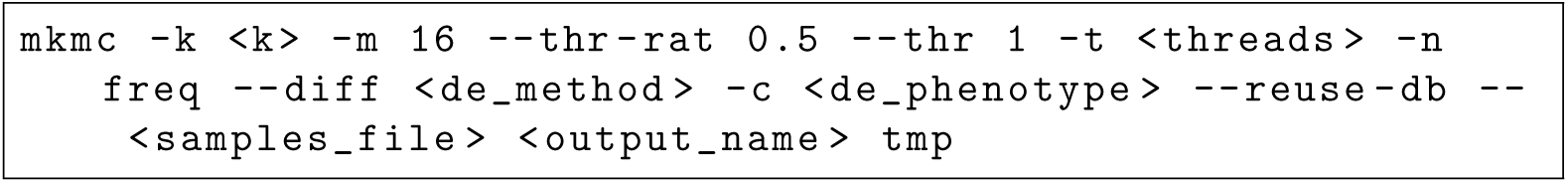

The binary matrix has been generated before the running. Normalization (-n freq) was not used for T-Test.

Additionally, ANOVA, WRS, and T-Test were run also with *p*-values normalization with a parameter --pval-corr b.

##### MKMC — postprocessing counts by computing entropy (d, e)

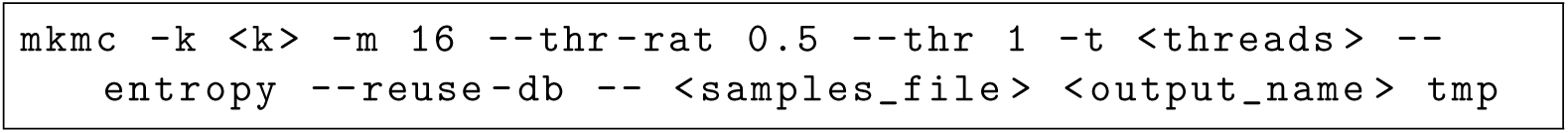

The binary matrix has been generated before the running.

##### MKMC — building a matrix and performing various tasks in single run (g; supplementary Fig. 9)

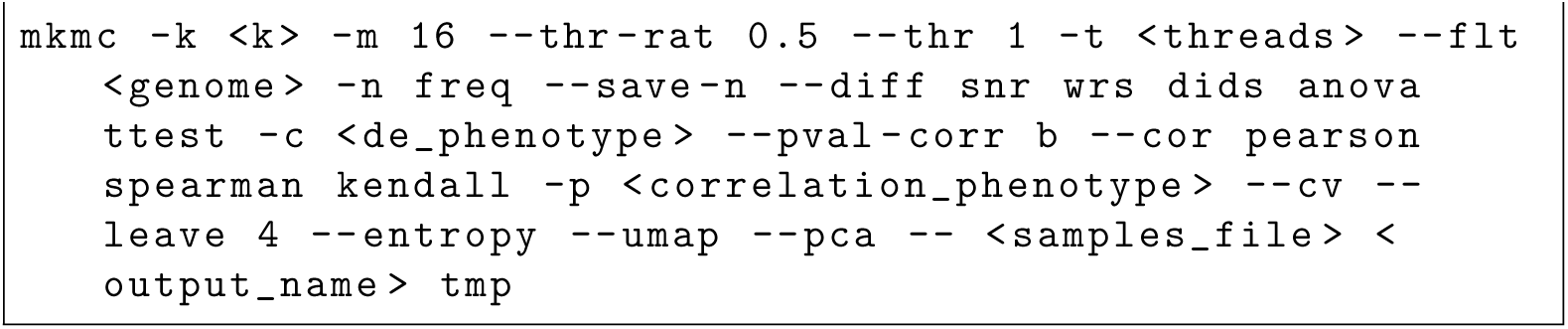

##### MKMC — building a matrix and performing various tasks in single run, but no dimensionality reduction (supplementary Fig. 10)

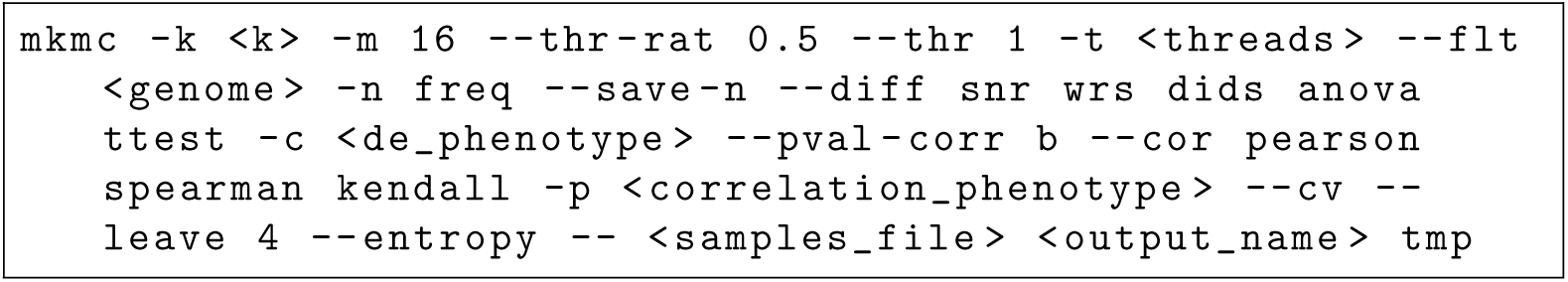

The binary matrix has not existed before the running.

Where:

- <k>— *k*-mer length,
- <threads> — number of threads,
- <correlation method> — one of pearson, spearman, kendall,
- <correlation phenotype> — a file with phenotype/design,
- <de method> — one of anova, dids, snr, ttest, wrs,
- <de phenotype> — a file with phenotype/design for differential *k*-mers analysis,
- <genome> — genome file (GCF 001465895.1 Nfu 20140520 genomic.fna) in FASTA format,
- <samples file> — a file with names of FASTQ files,
- <output file> — output files names template.

##### kmtricks — building a matrix (a, b, c, f, h)

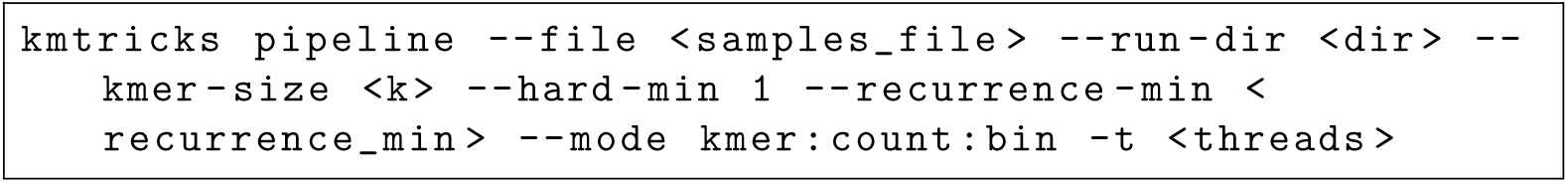

Where:

- <k> — *k*-mer length,
- <threads> — number of threads,
- <recurrence min> — a half of the number of samples,
- <samples file> — a file with names of FASTQ files,
- <dir> — an output directory.

##### Kamrat — building a matrix (a, b)

For every sample *i* = 1, 2*,…,N* Jellyfish was called:

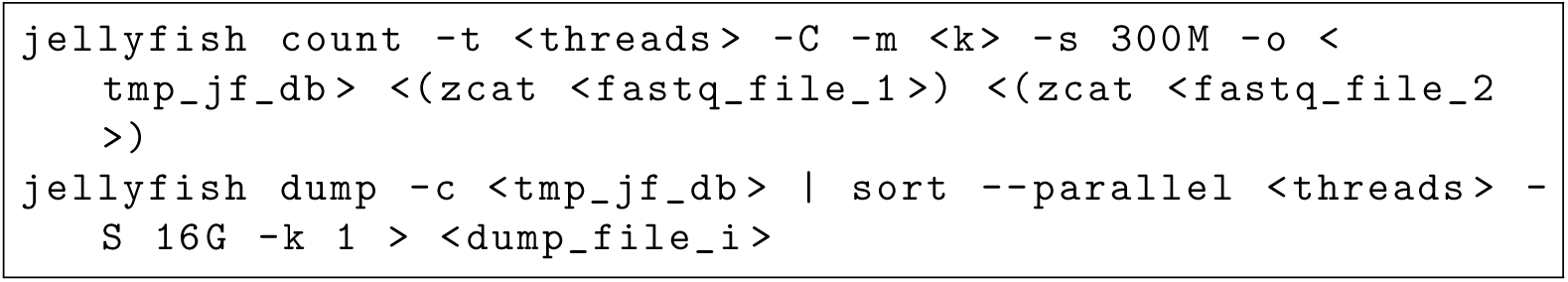

Then Jellyfish dumps were used to build a matrix in text form and run Kamrat:

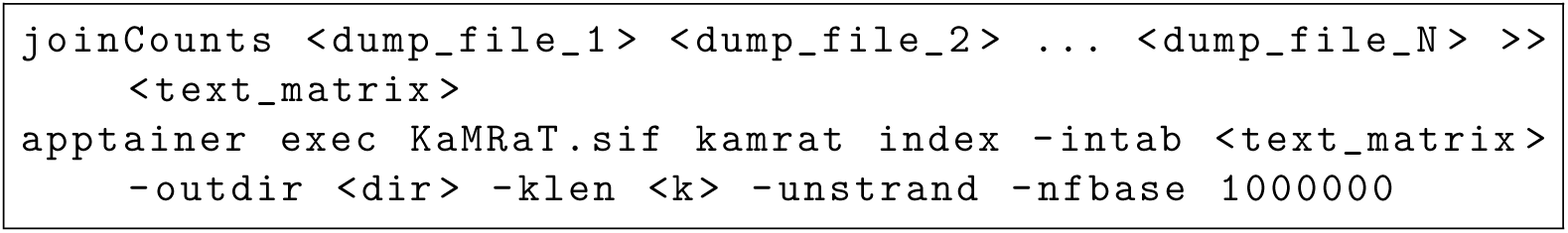

##### Kamrat — scoring k-mers (the matrix postprocessing) with design (d)

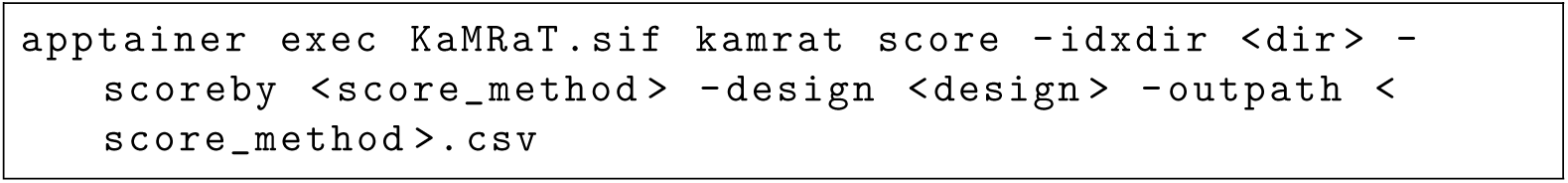

##### Kamrat — scoring k-mers (the matrix postprocessing) by computing entropy (d)

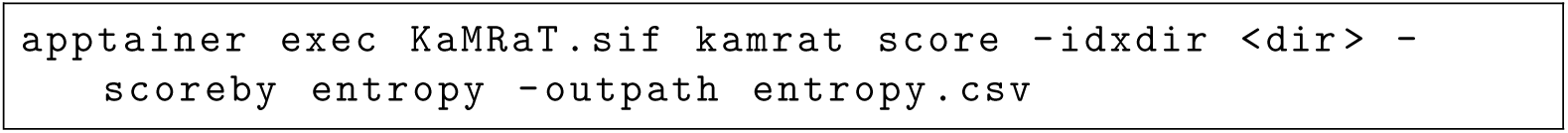

Where:

- <k> — *k*-mer length,
- <threads> — number of threads,
- <fastq file 1>, <fastq file 2> — paired input FASTQ files for single sample,
- <tmp_jf_db> — Jellyfish output database,
- <dump file 1>, <dump file 2>, <dump file i>, <dump file N> — joinCounts consecutive output files,
- <text matrix> — the matrix in text format,
- <dir> — an output directory,
- <score method> — one of pearson, spearman, snr, ttest.padj (enables also Benjamini-Hochberg correction), dids,

<design> — a file with phenotype/design.

## References

[1] Moeckel, C., Mareboina, M., Konnaris, M.A., Chan, C.S.Y., Mouratidis, I., Montgomery, A., Chantzi, N., Pavlopoulos, G.A., Georgakopoulos-Soares, I.: A survey of k-mer methods and applications in bioinformatics. Computational and Structural Biotechnology Journal 23, 2289–2303 (2024) 10.1016/j.csbj.2024.05.025

[2] Srivastava, A., Malik, L., Sarkar, H., Zakeri, M., Soneson, C., Love, M.I., Kingsford, C., Patro, R.: Alignment and mapping methodology influence transcript abundance estimation. Genome Biology 21(1), 239 (2020) 10.1186/s13059-020-02151-8

3. [3] Zhang, Q., Shao, M.: Transcript assembly and annotations: Bias and adjustment. PLoS Computational Biology 19(12), 1011734 (2023) 10.1371/journal.pcbi.1011734

[4] Zielezinski, A., Vinga, S., Almeida, J., Karlowski, W.M.: Alignment-free sequence comparison: benefits, applications, and tools. Genome Biology 18(1), 186 (2017) 10.1186/s13059-017-1319-7

[5] Sundaram, A., Tengs, T., Grimholt, U.: Issues with rna-seq analysis in non-model organisms: A salmonid example. Developmental & Comparative Immunology 75, 38–47 (2017) 10.1016/j.dci.2017.02.006. Impact of high throughput sequencing on comparative immunogenomics

[6] da Fonseca, R.R., Albrechtsen, A., Themudo, G.E., Ramos-Madrigal, J., Sibbesen, J.A., Maretty, L., Zepeda-Mendoza, M.L., Campos, P.F., Heller, R., Pereira, R.J.: Next-generation biology: Sequencing and data analysis approaches for non-model organisms. Marine Genomics 30, 3–13 (2016) 10.1016/j.margen.2016.04.012

[7] Xue, H., Gallopin, M., Marchet, C., Nguyen, H.N., Wang, Y., Lainé, A., Bessiere, C., Gautheret, D.: Kamrat: A c++ toolkit for k-mer count matrix dimension reduction. Bioinformatics 40(3), 090 (2024) 10.1093/bioinformatics/btae090

[8] Kukurba, K.R., Montgomery, S.B.: Rna sequencing and analysis. Cold Spring Harbor Protocols 2015(11), 951–969 (2015) 10.1101/pdb.top084970

[9] Conesa, A., Madrigal, P., Tarazona, S., et al.: A survey of best practices for rna-seq data analysis. Genome Biology 17, 13 (2016) 10.1186/s13059-016-0881-8

10. Dobin, A., Davis, C.A., Schlesinger, F., Drenkow, J., Zaleski, C., Jha, S., Batut, P., Chaisson, M., Gingeras, T.R.: Star: Ultrafast universal rna-seq aligner. Bioinformatics 29(1), 15–21 (2013) 10.1093/bioinformatics/bts635

[11] Liao, Y., Smyth, G.K., Shi, W.: featurecounts: An efficient general-purpose program for assigning sequence reads to genomic features. Bioinformatics 30(7), 923–930 (2014) 10.1093/bioinformatics/btt656

[12] Patro, R., Duggal, G., Love, M.I., Irizarry, R.A., Kingsford, C.: Salmon provides fast and bias-aware quantification of transcript expression. Nature Methods 14, 417–419 (2017) 10.1038/nmeth.4197

[13] Kokot, M., Dl-ugosz, M., Deorowicz, S.: KMC 3: counting and manipulating k-mer statistics. Bioinformatics 33(17), 2759–2761 (2017) 10.1093/bioinformatics/btx304

[14] Deorowicz, S., Agnieszka, D., Kokot, M.: VCFShark: how to squeeze a vcf file. Bioinformatics 37(19), 3358–3360 (2021) 10.1093/bioinformatics/btab211

[15] Lemane, T., Medvedev, P., Chikhi, R., Peterlongo, P.: kmtricks: Efficient and flexible construction of bloom filters for large sequencing data collections. Bioinformatics Advances 2(1), 029 (2022) 10.1093/bioadv/vbac029

[16] McKay, A., Costa, E.K., Chen, J., Hu, C.-K., Chen, X., Bedbrook, C.N., Khondker, R.C., Thielvoldt, M., Singh, P.P., Wyss-Coray, T., Brunet, A.: An automated feeding system for the african killifish reveals the impact of diet on lifespan and allows scalable assessment of associative learning. eLife 11, 69008 (2022) 10.7554/eLife.69008

[17] Schaum, N., Lehallier, B., Hahn, O.e.a.: Aging hallmarks exhibit organ-specific temporal signatures. Nature 583, 596–602 (2020) 10.1038/s41586-020-2499-y

[18] Marçais, G., Kingsford, C.: A fast, lock-free approach for efficient parallel counting of occurrences of k-mers. Bioinformatics 27(6), 764–770 (2011) 10.1093/bioinformatics/btr011

[19] Audoux, J., Philippe, N., Chikhi, R., Salson, M., Gallopin, M., Gabriel, M., Le Coz, J., Drouineau, E., Commes, T., Gautheret, D.: DE-kupl: exhaustive capture of biological variation in RNA-seq data through k-mer decomposition. Genome biology 18(1), 243 (2017) 10.1186/s13059-017-1372-2

[20] Greenacre, M., Groenen, P.J.F., Hastie, T., Mazet, O., Meinshausen, N., Buja, A.: Principal component analysis. Nature Reviews Methods Primers 2, 100 (2022) 10.1038/s43586-022-00184-w

[21] McInnes, L., Healy, J., Melville, J.: Umap: Uniform manifold approximation and projection for dimension reduction. arXiv preprint arXiv:1802.03426 (2018) 10.48550/arXiv.1802.03426

[22] Liao, Y., Smyth, G.K., Shi, W.: featurecounts: an efficient general-purpose program for assigning sequence reads to genomic features. Bioinformatics 30, 923–930 (2014) 10.1093/bioinformatics/btt656

[23] Love, M.I., Huber, W., Anders, S.: Moderated estimation of fold change and dispersion for rna-seq data with deseq2. Genome Biology 15(12), 550 (2014) 10.1186/s13059-014-0550-8

[24] Law, C.W., Chen, Y., Shi, W., Smyth, G.K.: voom: precision weights unlock linear model analysis tools for rna-seq read counts. Genome Biology 15, 29 (2014) 10.1186/gb-2014-15-2-r29

[25] Li, Y., Ge, X., Peng, F., Li, W., Li, J.J.: Exaggerated false positives by popular differential expression methods when analyzing human population samples. Genome Biology 23, 79 (2022) 10.1186/s13059-022-02648-4

[26] Choi, H.M.T., Schwarzkopf, M., Fornace, M.E., Acharya, A., Artavanis, G., Stegmaier, J., Cunha, A., Pierce, N.A.: Third-generation in situ hybridization chain reaction: multiplexed, quantitative, sensitive, versatile, robust. Development 145(12), 165753 (2018) 10.1242/dev.165753

[27] Brooks, T.G., Lahens, N.F., Mrčela, A., et al.: Challenges and best practices in omics benchmarking. Nature Reviews Genetics 25, 326–339 (2024) 10.1038/s41576-023-00679-6

[28] Li, B., Dewey, C.N.: Rsem: Accurate transcript quantification from rna-seq data with or without a reference genome. BMC Bioinformatics 12, 323 (2011) 10.1186/1471-2105-12-323

[29] Bray, N.L., Pimentel, H., Melsted, P., Pachter, L.: Near-optimal probabilistic rna-seq quantification. Nature Biotechnology 34, 525–527 (2016) 10.1038/nbt.3519

[30] Lemane, T., Chikhi, R., Peterlongo, P.: Kmdiff, large-scale and user-friendly differential k-mer analyses. Bioinformatics 38(24), 5443–5445 (2022) 10.1093/bioinformatics/btac645

[31] Marçais, G., Kingsford, C.: A fast, lock-free approach for efficient parallel counting of occurrences of k-mers (2011). 10.1093/bioinformatics/btr011. https://academic.oup.com/bioinformatics/article/27/6/764/234905

[32] Sacomoto, G.A., Kielbassa, J., Chikhi, R., Uricaru, R., Antoniou, P., Sagot, M.-F., Peterlongo, P., Lacroix, V.: Kissplice: De-novo calling alternative splicing events from rna-seq data. BMC Bioinformatics 13(S6), 5 (2012) 10.1186/1471-2105-13-S6-S5

[33] Drouin, A., Giguère, S., Dèraspe, M., Marchand, M., Tyers, M., Loo, V.G., Bourgault, A.-M., Laviolette, F., Corbeil, J.: Predictive computational phenotyping and biomarker discovery using reference-free genome comparisons. BMC Genomics 17, 530 (2016) 10.1186/s12864-016-2846-6

[34] Audoux, J., Philippe, N., Chikhi, R., Salson, M., Gallopin, M., Gabriel, M., Le Coz, J., Drouineau, E., Commes, T., Gautheret, D.: De-kupl: Exhaustive capture of biological variation in rna-seq data through k-mer decomposition. Genome Biology 18, 141 (2017) 10.1186/s13059-017-1372-2

[35] Rahman, A., Hallgrímsdóttir, I., Eisen, M., Pachter, L.: Association mapping from sequencing reads using k-mers. eLife 7, 32920 (2018) 10.7554/eLife.32920

[36] Thomas, A., Barriere, S., Broseus, L., Brooke, J., Lorenzi, C., Villemin, J.-P., Beurier, G., Sabatier, R., Reynes, C., Mancheron, A., al.: Gecko is a genetic algorithm to classify and explore high-throughput sequencing data. Communications Biology 2, 246 (2019) 10.1038/s42003-019-0456-9

[37] Lorenzi, C., Barriere, S., Villemin, J.-P., Bretones, L.D., Mancheron, A., Ritchie, W.: Imoka: K-mer based software to analyze large collections of sequencing data. Genome Biology 21, 292 (2020) 10.1186/s13059-020-02165-2

[38] Tayyebi, Z., Pine, A.R., Leslie, C.S.: Scalable and unbiased sequence-informed embedding of single-cell atac-seq data with cellspace. Nature Methods 21, 1014–1022 (2024) 10.1038/s41592-024-02274-x

[39] Chaung, K., Baharav, T., Henderson, G., et al.: Splash: A statistical, reference-free genomic algorithm unifies biological discovery. Cell 186(24), 5440–545626 (2023) 10.1016/j.cell.2023.10.026

[40] Kokot, M., Dehghannasiri, R., Baharav, T., et al.: Scalable and unsupervised discovery from raw sequencing reads using splash2. Nature Biotechnology 43, 1084–1090 (2025) 10.1038/s41587-024-02381-2

[41] Dehghannasiri, R., Kokot, M., Starr, A.L., Maziarz, J., Gordon, T., Tan, S.Y., Wang, P.L., Voskoboynik, A., Musser, J.M., Deorowicz, S., Salzman, J.: sc-splash provides ultra-efficient reference-free discovery in barcoded single-cell sequencing. bioRxiv (2024) 10.1101/2024.12.24. Preprint

[42] Chen, J., Khondker, R.C., Brunet, A.: Breeding and reproduction of the african turquoise killifish nothobranchius furzeri. Cold Spring Harbor Protocols 2023(6), 107816 (2023) 10.1101/pdb.prot107816

[43] Costa, E.K., Chen, J., Guldner, I.H., Mboning, L., Schmahl, N., Tsenter, A., Wu, M.-R., Moran-Losada, P., Bouchard, L.S., Wang, S., Singh, P.P., Pellegrini, M., Brunet, A., Wyss-Coray, T.: Multi-tissue transcriptomic aging atlas reveals predictive aging biomarkers in the killifish. bioRxiv (2025) 10.1101/2025.01.28.635350 https://www.biorxiv.org/content/early/2025/02/01/2025.01.28.635350.full.pdf

[44] Schindelin, J., Arganda-Carreras, I., Frise, E., et al.: Fiji: an open-source platform for biological-image analysis. Nature Methods 9(7), 676–682 (2012) 10.1038/nmeth.2019

[45] Golub, T.R., Slonim, D.K., Tamayo, P., Huard, C., Gaasenbeek, M., Mesirov, J.P., Coller, H., Loh, M.L., Downing, J.R., Caligiuri, M.A., et al.: Molecular classification of cancer: class discovery and class prediction by gene expression monitoring. Science 286(5439), 531–537 (1999)

[46] Ronde, J., Rigaill, G., Rottenberg, S., Rodenhuis, S., Wessels, L.: Identifying subgroup markers in heterogeneous populations. Nucleic Acids Research 41(21), 200–200 (2013)

